# A Hybrid Physics-Deep Learning Framework for Combinatorial De Novo Design of Small-Molecule Binding Proteins

**DOI:** 10.64898/2026.04.12.717919

**Authors:** Connor V. Galvin, Amy B. Guo, Maple N. Chen, Isabella L. Alfonso, Dominic Grisingher, Simon Kretschmer, Divya Kranthi, Mark J. S. Kelly, Tanja Kortemme

## Abstract

Engineering small-molecule binding proteins de novo remains a significant challenge as even advanced generative models struggle to model the atom-level details of protein-ligand interactions with sufficient accuracy. Higher experimental success rates have resulted from methods that explicitly scaffold predefined binding interactions into helical bundles. Here we introduce a scaffolding strategy that generalizes to alpha-beta architectures. By screening thousands of combinatorially assembled protein-ligand interactions against diverse de novo backbones with finely varied pocket geometries, the protocol allows for high-fidelity accommodation of target interaction geometries. Our protocol then integrates physics-based and deep learning methods for optimization of interfacial interactions and sequence-structure compatibility, considerably improving *in silico* design metrics. Applying this method to two chemically similar steroids achieved a notable experimental success rate (4/26 designs bind their targets), and NMR structures of two designs are in good agreement with design models. Our generalizable, atomically precise approach offers a robust framework for small-molecule binder design, effectively eliminating the need for high-throughput screening.

## INTRODUCTION

Small molecule binding is a fundamental ability of natural proteins, mediating numerous processes from signaling to metabolism. Consequently, the ability to engineer new proteins to bind desired target molecules has enormous technological utility (*1*) in applications such as biosensing (*2*) (*3*) (*4*), drug binding (*5*) and delivery, and catalysis (*6*, *7*). Computational protein design offers the prospect of on-demand production of small molecule binders, with control over critical parameters such as selectivity and binding-induced response. However, although there have been successes in the field of *de novo* design of small molecule binders (*4*) (*5*) (*8*) (*9*), a reliable and generalizable strategy for producing *de novo* proteins binding with high affinity to any desired target, in particular those that are highly polar and flexible, with success rates substantially above 1%, has not yet been realized.

A key difficulty in the *de novo* design of small molecule binding proteins lies in satisfying the dual requirements of designing both the atomically detailed interactions between the protein and target molecule, along with the overall protein topology and ligand orientation (*10*); considering these degrees of freedom simultaneously with the required accuracy remains challenging (all atom co-generation models (*11*, *12*) are capable of this in principle, but have not yet produced high-affinity binding proteins for diverse targets at high success rates). Existing approaches to the design of small molecule binders can thus be classified based on the order in which these elements - designing an overall scaffold with a binding site and accommodating atomically detailed interactions with the ligand in that binding site - are addressed.

Most generative machine learning models for protein design operate by first producing a protein backbone and relative orientation of a binding partner (*13*). In subsequent steps design algorithms (*14*) are applied to identify sequences which can both stabilize the backbone and satisfy the detailed interactions with the binding partner. While this workflow has proven remarkably effective at producing well folded proteins and protein-protein binders, hydrogen bonding interactions critical for small molecule binding are often exquisitely constrained in their interaction geometries (*15*) and have energetic tradeoffs that are difficult to model accurately. For example, the desolvation penalty incurred by the burial of polar ligand atoms within a binding pocket must be compensated by the formation of new hydrogen bonds with the protein, and these are more difficult to satisfy than the hydrophobic packing interactions that stabilize a protein core or a larger protein-protein interface. The result is low experimental success rates for small molecule binders generated with diffusion methods (<0.1%) (*15*), necessitating high throughput screening of large numbers of designs to identify functional binders, which is not always possible for a given target.

Alternatively, a set of key interactions between protein and target molecule (a binding ‘motif’) can be explicitly defined up front, and then built into a protein backbone (‘scaffold’) that can accommodate the motif and ligand with the desired geometries (*16*). The main limitation of this approach is the low probability of compatibility between a given motif-scaffold pair (*17*) (on the order of 10^-4^ for a small 3 residue motif and rapidly decreasing with increasing number of motif residues). While diffusion methods have also been used for the problem of “motif scaffolding” (*17*) - generating a new backbone around a predefined motif - these approaches often produce either distorted structures that accommodate the motif but are not designable, or regular structures that do not accommodate the motif precisely enough. Helical parametrization (*18*) (*19*) can be used to get around these problems because it allows one to generate thousands of backbones with tunable variation and thus increases the likelihood to accommodate pre-defined constellations of discrete amino acid geometries with high fidelity (*19*). However, this method is limited to the creation of helical bundles, which in turn limits the scope of the target molecules that can be accommodated within the typical bundle core and the range of functions that can be achieved.

Here, we present an approach for precisely scaffolding protein-ligand interactions that generalizes to alpha-beta topologies typical in natural small-molecule binding proteins. We combine computational methods that generate (i) thousands of diverse binding motifs (*20*) (**Figure 1A**), and (ii) thousands of finely tunable protein backbones (**Figure 1B**) into which these motifs can be placed (*21*) (**Figure1C**). We show that together, these methods allow for high fidelity scaffolding of idealized binding interactions for a variety of target molecules. Further, we develop a complete workflow for optimizing the designed proteins for stability and binding, using ProteinMPNN (*22*) and Rosetta for sequence design, Rosetta FastRelax for minimization and filtering, AlphaFold2 (*23*) for evaluating designability, and a custom Rosetta-based algorithm for the refinement of hydrogen bonding features within the binding pocket. We show that this workflow, termed CLAIRE (<u>C</u>ombinatoria<u>L A</u>ssembly with Integrated <u>RE</u>finement), which iterates between machine-learning and physics-based methods (**Figure 1A-C**), is effective for generating functional binders with a high experimental success rate amenable to small scale expression and screening, using as targets the steroids progesterone and estriol.

**Figure 1:**
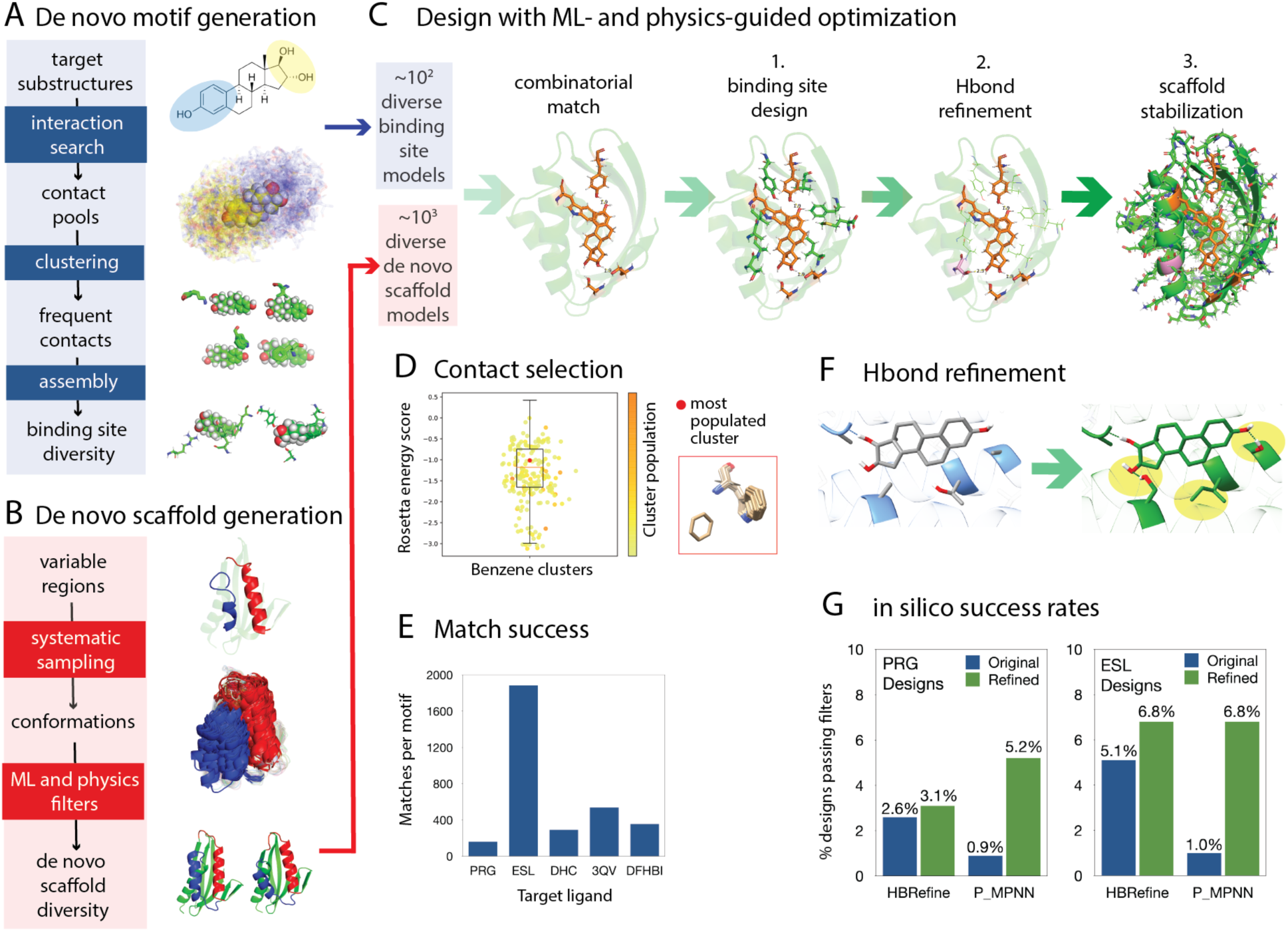
CLAIRE Design workflow and improved success rates from refinement steps. **A**) Motif generation protocol. Target ligand substructures are selected, and all interactions between those substructures and protein side chains are obtained from the PDB. The resultant contact pools are clustered and combinatorially assembled into unique binding motifs favoring the most highly represented interactions. **B**) De novo scaffold generation protocol. Variable regions in an input de novo NTF2 protein (PDB 5TPJ) are systematically sampled. The resulting conformations are filtered based on both deep learning and physics-based criteria, resulting in a diverse scaffold library with high predicted stability. **C**) The motif and scaffold libraries are combinatorially screened for compatibility. The sequences of the initial matches are optimized for 1. binding to the target, 2. hydrogen bond satisfaction in the binding site, and 3. stability. **D**) Box and whisker plot showing distribution of Rosetta energy scores across clustered interactions with a benzene ring, colored by cluster population (left). The most highly populated (but not best scoring) cluster for benzene is an edge to face pi stacking interaction with a tryptophan (right). **E**) Number of buried matches obtained for five diverse small molecules with a range of sizes and chemical groups screened against 1,816 de novo NTF2 scaffolds (Methods). **F**) Example binding site before (left) and after (right) HBRefine algorithm. Two additional hydrogen bonds have been formed with the unsatisfied oxygen atoms of the target small molecule (estriol), and one buried threonine with an unsatisfied oxygen has been mutated to isoleucine (yellow shading). **G**) Improved in silico success rates from design refinement methods, HBRefine for binding site polar features and ProteinMPNN for scaffold stabilization, for designed progesterone (left) and estriol (right) binders (see **Supplementary Table S7**).

## RESULTS

Our overall strategy involved to generate, first, diverse binding site motifs and second, diverse scaffolds into which these motifs could be placed. For the generation of small molecule binding motifs (**Figure 1A**), we used the Binding Sites from Fragments (BSFF) method (*20*). BSFF first decomposes target molecules into smaller chemical substructures and then scrapes the PDB for all interacting amino acids. We filtered the resultant interaction pools to eliminate duplicates or high sequence identity (>70%) entries.

Improvements in the design of stable protein folds came from leveraging the wealth of structural data in the PDB, allowing for the derivation of sequence and structural preferences that can be used to both considerably narrow the sample space and improve scoring functions (*24*). We therefore hypothesized that binding site design could similarly benefit from an approach that considers common interaction modes between protein side chains and small molecule substructures. Accordingly, we spatially clustered the amino acid side chains interacting with a given small molecule substructure into discrete interaction modes and retained those that were statistically overrepresented. We observed strong statistical preferences for particular interaction modes with each chemical substructure we queried, with on average 1% of clusters accounting for 15% of all interactions (**Supplementary Figure S1**). Further, the Rosetta energy scores (*25*) of these interactions were found not to correlate with their observed frequencies. In particular, the scores did not reflect the strong observed preferences for pi stacking (**Figure 1D**) and cation-pi interactions, as well as cases with higher order hydrogen bonding (one amino acid residue hydrogen bonding simultaneously with multiple ligand polar atoms). Finally, 4-residue combinations of interactions were enumerated combinatorially and those without steric clashes were added to the motif library used for design (**Figure 1A**).

For the generation of geometrically diverse scaffold proteins (**Figure 1B**), we used the Loop-helix-loop Unit Combinatorial Sampling (LUCS) method (*21*). LUCS operates by systematically varying the geometries (i.e., positions, lengths and orientations) of helices in an input protein. This strategy mimics how nature repurposes a single protein fold to accommodate a variety of functions by finely varying the geometries of structural elements about the functional site, while also producing novel geometries not seen in the ensemble of naturally occurring examples for the input protein fold. We used LUCS to reshape *de novo* proteins with nuclear transport factor 2 (NTF2) topology (*26*), as this fold was previously shown to be amenable to LUCS reshaping (40% of designs that were tested were well folded (*21*)) and frequently binds small molecules in nature, with a cone like structure containing a large central cavity between several helices capping an underlying curved beta-sheet.

To begin, we tested whether our design strategy would be able to scaffold binding motifs for a range of small molecules with a diversity of sizes and chemical groups, applying it to 5 molecules ranging from 180 to 314 daltons: progesterone (PRG), estriol (ESL), caffeic acid (DHC), 3-cyano-7-hydroxy coumarin (3QV), and 3,5-difluoro-4-hydroxybenzylidene imidazolinone (DFHBI). For each molecule, we generated interaction motifs using BSFF as described above. We then used the Rosetta match application (*16*) to graft the binding motifs into 1,816 LUCS generated NTF2 scaffolds. The algorithm functions by building individual motif interactions at user-defined candidate positions within the scaffold protein. The interaction geometries are characterized by six degrees of freedom, which in our case were narrowly bounded, with distances held within 0.5 Å and all angles and torsions held within 10-15° of ideal values (**Supplementary Figure S2**). The algorithm checks that all geometric constraints are satisfied and that there are no clashes with the protein backbone, and then bins the resultant structure based on the protein-ligand orientation. A ‘match’ is obtained when all motif interactions can be built into a given scaffold with the relative ligand orientations occupying the same bin. We then filtered the resulting grafted structures to ensure sufficient burial of the target molecules within the binding pocket, discarding those with greater than 30% of the ligand surface area solvent exposed. The matching success rate is shown in **Figure 1E**, with greater than 160 ‘matches’ per input motif obtained in all cases, which is sufficient to produce >10,000 grafted structures from 100 motifs with our set of 1,816 NTF2 scaffolds.

To develop the next steps of our design and optimization workflow and ultimately validate designs experimentally, we selected the target molecules progesterone and estriol. These molecules are largely isosteric steroids, with the primary exception being that progesterone has two carbonyl groups at the 3 and 17 positions, while estriol has two hydroxyl groups at these same positions, along with one additional hydroxyl group at position 16 (**Supplementary Figure S3**). Adequate satisfaction of ligand polar atoms is one of the primary challenges in small molecule binder design, due to the greater geometric constraints for hydrogen bonding relative to hydrophobic packing, requiring increased precision for the relevant interacting residues in the designed protein. We reasoned that these molecules would provide an interesting case for testing the effect of one additional ligand polar atom on design success.

We selected 8,500 buried matches for each of estriol and progesterone for design. Our design workflow proceeded through three steps of sequence and structure optimization, scoring, and filtering (**Figure 1C**). In the first step (binding site design), we used the Rosetta FastDesign algorithm to design all binding site residues (all side chains with at least one heavy atom within 5 Å of ligand) in the matches, with protein-ligand interactions up-weighted by 2.5-fold relative to protein-protein interactions. We then filtered these designs by binding site quality criteria (**Methods**), yielding 432 and 219 designs for estriol and progesterone, respectively.

When analyzing the resulting designs, we noticed a considerable number of buried unsatisfied hydrogen bonding donors and acceptors on both the ligand and the protein residues within the binding site. We therefore developed a custom binding site hydrogen bond refinement method, HBRefine, and applied it to these designs. The method introduces mutations to form new hydrogen bonds between the protein and any unsatisfied ligand polar atoms. It also identifies protein residues with buried unsatisfied polar atoms and mutates them to favorable non-polar residues (**Figure 1F**). The resultant designs were filtered to ensure that there were no remaining buried unsatisfied polar atoms in the binding site. Additional filtering criteria included protein-ligand shape complementarity, the Rosetta ‘contact molecular surface’ metric (a measure of the Lawrence and Coleman shape complementarity between the protein and ligand surfaces with penalties applied for cavities that are smaller than the spherical probe used for surface identification), binding site preorganization (Rosetta RotamerBoltzmann metric, an average of the approximated Boltzmann probabilities for binding site rotamers), and binding energy (Rosetta ddG metric, the calculated difference between the Rosetta energy of the complex and the individual components). These criteria were derived from a set of crystal structures of monomeric protein-small molecule complexes with experimentally determined affinity values (*27*). The average values for relevant metrics across these known binders (**Supplementary Figure S4)** were used to inform the filtering thresholds for our designs (**Supplementary Table S1**).

Next, we used ProteinMPNN (*22*) to redesign all residues outside of the binding site, and used the resultant sequence profiles as input to the Rosetta FastDesign algorithm to redesign these residues within the physical context of the binding site. We then filtered these designs by all previously evaluated binding centric metrics, as well as additional criteria relating to protein stability, such as high packing efficiency (Rosetta ‘packstat’ metric, which compares per residue van der Waals scores with averages for high quality crystal structures), exposed hydrophobic surface area (Rosetta ExposedHydrophobics filter, calculates SASA for all hydrophobic residues), and a lack of either unsatisfied or oversaturated polar atoms across the protein. We found that these targeted refinement steps implemented in our design workflow, HBrefine and ProteinMPNN, increased *in silico* success rates up to ∼7-fold (**Figure 1G**).

To compare our overall workflow, CLAIRE, with another recent design method, we also used RosettaFold Diffusion all-atom (*22*) to produce designs for estriol and progesterone following the published strategy employed for the steroid digoxigenin. We subjected ∼30,000 diffused binders for each target (see **Methods**) to our filtering criteria and compared the results with those of CLAIRE designs. Our designs achieved considerably higher *in silico* success rates (satisfaction of all quality criteria) for both targets (**Figure 2A**), with the main point of failure for the diffused designs being insufficient hydrogen bonds to the ligand and unsatisfied hydrogen bond donors and acceptors buried in the binding site (**Figure 2B**). Additionally, the diffused designs are almost exclusively helical bundle proteins, compared with the alpha-beta topology of our designs (**Supplementary Figure S5**). Across the other quality criteria, the diffused designs performed comparably to those from CLAIRE, with the exception of a higher fraction of designs passing protein-ligand shape complementarity and packing filters (**Figure 2B, Supplementary Figure S6**).

**Figure 2:**
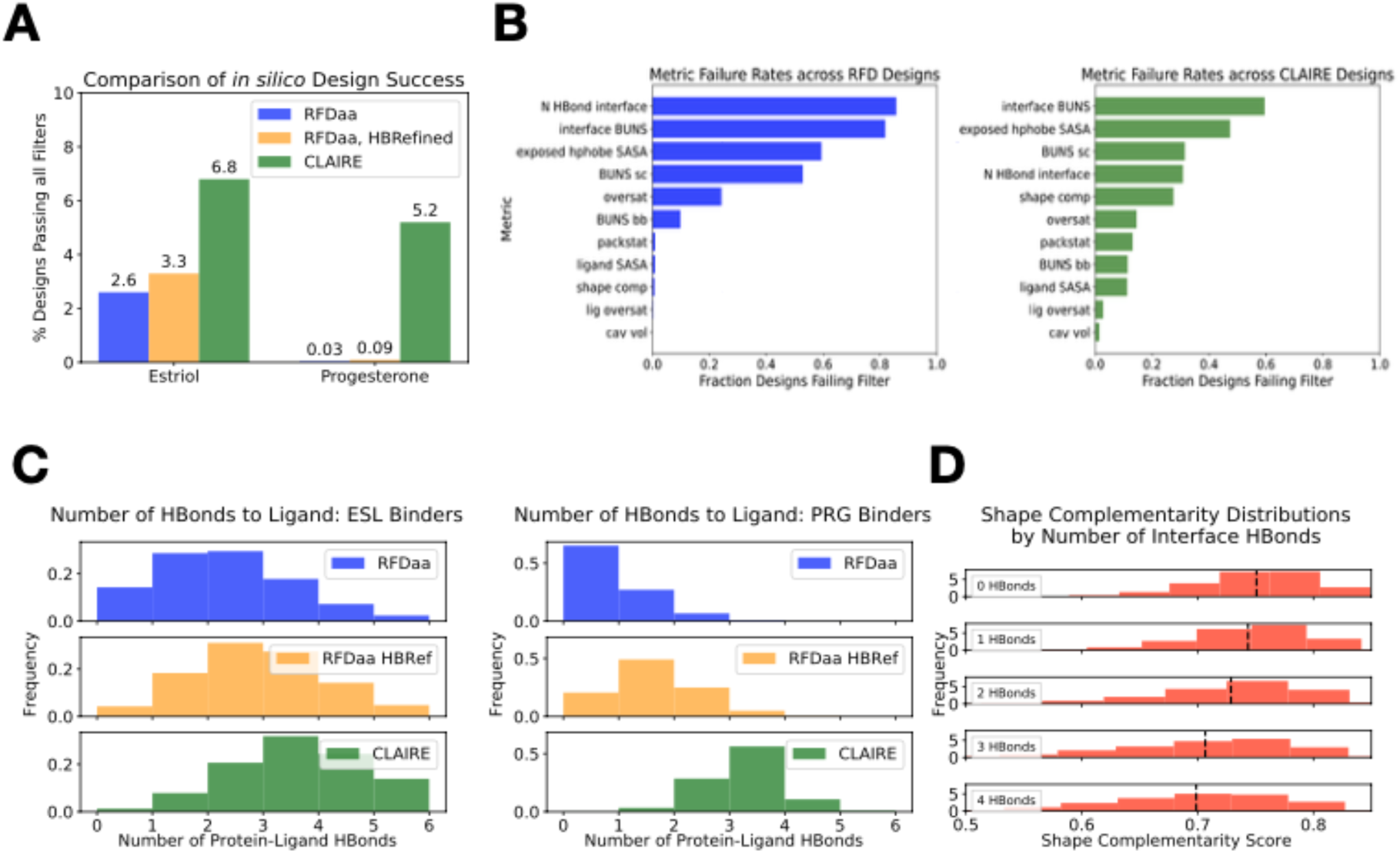
Comparison between designed progesterone and estriol binders from RosettaFold Diffusion all-atom (RFDaa) and CLAIRE. **A**) Overall *in silico* success rates for progesterone and estriol binders for the RFDaa designs, the RFDaa designs after refinement by our HBRefine method, and CLAIRE designs. While HBRefine can rescue a number of RFDaa designs, CLAIRE designs still have higher *in silico* success rates overall. **B**) Failure rates for individual filtering criteria across RFDaa designs (left) and CLAIRE designs (right). Protein-ligand hydrogen bond satisfaction (N HBond interface=number protein-ligand hydrogen bonds, cutoff 3; interface BUNS=number of buried unsatisfied polar atoms at the binding interface, cutoff 0) is the largest cause of failure for RFDaa designs. Details on all filters can be found in **Supplementary Table S1**. **C**) Histograms showing distributions for the number of protein-ligand hydrogen bonds across designs. HBRefine shifts the distributions for RFDaa designs towards higher values, but designs generated with CLAIRE still have higher prevalence of interface hydrogen bonding. **D**) The distributions of Rosetta protein-ligand shape complementarity values across all designs, for designs with different numbers of interface hydrogen bonds. Shape complementarity appears to decrease with increased hydrogen bonding, suggesting a tradeoff between these two features.

To see whether the hydrogen bonding features of the diffused designs could be improved, we subjected them to our targeted hydrogen bond refinement method. While a substantial number of the diffused designs could be rescued in this manner (**Figure 2C)**, the overall success rates remained lower than for the designs from our method (3.3% compared with 6.8% for estriol, 0.09% compared with 5.2% for progesterone (**Figure 2A)**. This result is not unexpected, as our method benefits from the ability to explicitly define hydrogen bonding interactions with all ligand polar atoms up front, while diffusion relies on the prospect of designing in these interactions after establishing a protein-ligand orientation. Our results suggest that satisfying hydrogen bonding geometries with the required precision and completeness at later stages of the design process is often unachievable. Instead, it appears that there exists a trade-off between protein-ligand shape complementarity and the ability to satisfy the geometric constraints of intermolecular hydrogen bonding (**Figure 2D**).

We proceeded with applying our CLAIRE design workflow developed above to large-scale production runs for experimental testing. After subjecting our full set of buried estriol and progesterone matches (52,862 for estriol, 64,238 for progesterone, **Figure 3A, Methods**) to binding site design (step 1 in **Figure 1C**), hydrogen bond and stability refinement (steps 2 and 3 in **Figure 1C**) and stringent physics-based filtering, we retained 1,702 and 745 designs for estriol and progesterone, respectively. We note that using naturally existing binding site motifs (2 for estriol and 16 for progesterone) and matching them into 24 different single domain NTF2 folds with available experimental structures did not result in any designs passing our first round of binding site quality filters (**Figure 3A**), highlighting the advantage of our methods for *de novo* generation of both motifs (**Figure 1A**) and scaffolds (**Figure 1B**). Finally, we used Alphafold2 (*23*) to predict structures for our designed sequences, and filtered for high confidence (average pLDDT ≥ 88.0) and low RMSD between the input and predicted structures (Ca RMSD ≤ 1.5 Å). Thirteen designs for each target passing these filters were ordered for experimental characterization (**Supplementary Tables S2, S3**). The structures of these 26 designs were also well predicted with the recently published Boltz-2 (*28*) (although it failed to recapitulate the designed small molecule binding pose, **Supplementary Tables S4, S5, Supplementary Figure S7**).

**Figure 3:**
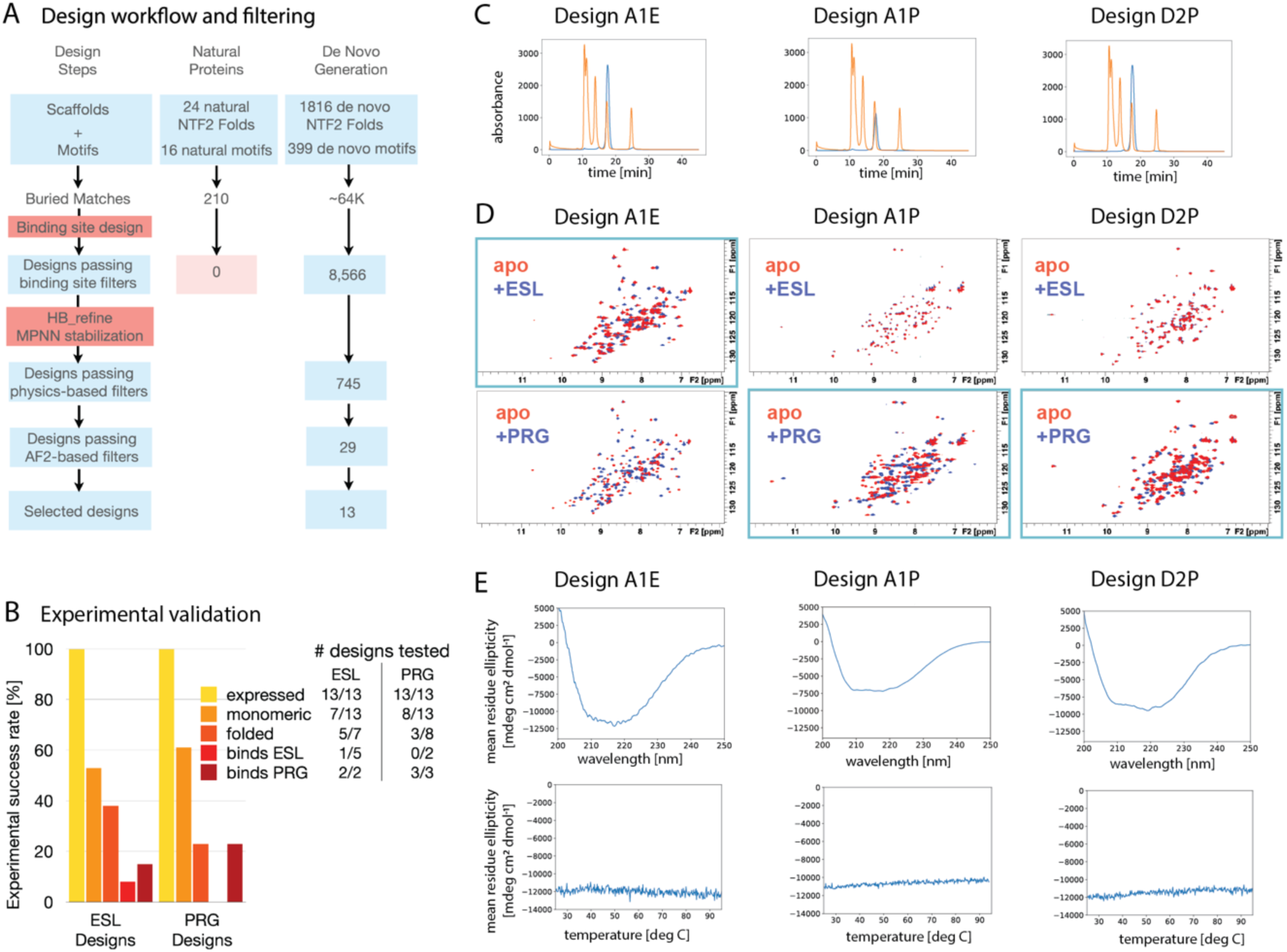
Experimental validation of designed estriol and progesterone binders. **A**) Design approach showing numbers of designs at each step for progesterone binders (for estriol binder design see **Methods**). Using natural NTF2 folds and progesterone binding motifs did not yield any designs passing initial binding site quality filters. **B**) Summary of experimental success rates with numbers of characterized designs; EST: estriol; PRG: progesterone **C-E)** Biophysical characterization of 3 designs (designed ESL binder A1E left, designed PRG binders A1P middle, D2P right). **C)** SEC traces showing the designs to be monomeric. **D**) ^15^N NMR HSQC chemical shift perturbation (CSP) data with the spectra for apo proteins shown in blue and the spectra for holo protein with ligands shown in red. Top row shows CSP upon addition of ESL, bottom row upon addition of PRG. On-target combinations of designs are highlighted with light blue box. A1E binds both on- and off-target ligands, while the two designs A1P and A2P are specific to their target PRG. **E**) CD spectra (top) and CD thermal melt monitored at 222 nm (bottom) shows the three designs have well defined secondary structure, and are highly thermostable up to 95 C.

All 26 designs expressed in the soluble fraction of *E. coli* and were evaluated for oligomeric states, folding, and stability using size exclusion chromatography (SEC) and nuclear magnetic resonance (NMR) spectroscopy (**Figure 3B, Supplementary Table S6**). In total, 15/26 (∼58%) of the tested designs were monomeric according to SEC (**Figure 3C**, **Supplementary Figures S8, S9**), and 8/26 (31%) of the tested designs were well folded according to ^15^N Heteronuclear Single Quantum Coherence (HSQC) NMR spectra. This set included 5 of the estriol binder designs and 3 of the progesterone binder designs (**Figure 3B**).

We then screened for binding activity using chemical shift perturbation experiments for the ^15^N HSQC spectra of the designs in the presence and absence of ligand (**Figure 3D, Supplementary Figures S10, S11**). One of the designed estriol binders (A1E) showed chemical shift perturbations in the presence of estriol, and 3 of the designed progesterone binders (A1P, C2P, D2P) showed chemical shift perturbations in the presence of progesterone, equating to success rates of 7.6% and 23.1%, respectively (**Figure 3B, Supplementary Figures S10, S11 Supplementary Table S6**). In the same manner, we tested the specificity of these designs, with two of the designed estriol binders showing binding to progesterone, while neither of the tested progesterone binders showed a clear binding signal with estriol under the conditions tested (**Supplementary Figure S12**). The greater promiscuity of the estriol binder designs towards progesterone can be rationalized by the compatibility between the protein-estriol hydrogen bonds and the carbonyl oxygens of progesterone, which are nearly isosteric with the equivalent hydroxyl groups in estriol (**Supplementary Figure S13**). While the progesterone binder designs can in a similar manner hydrogen bond with two of the hydroxyl groups in estriol, they have no residues designed to satisfy the additional hydroxyl oxygen present in estriol.

We selected three designs with the most pronounced ligand-induced chemical shift perturbations, one for estriol (A1E) and two for progesterone (A1P, D2P), for additional experimental characterization. Thermal melts for these three proteins revealed them to be highly thermostable, with none showing changes in the CD signal at 222 nm up to 95 deg C (**Figure 3E)**. We attempted to quantify the binding affinity of these designs for their target molecules using Isothermal Titration Calorimetry but were unsuccessful due to low signal-to-noise and high heats of dilution in the DMSO cosolvent conditions required for dissolving the ligands. Instead, we performed a ligand titration for D2P monitored by 2D ^15^N HSQC NMR spectra. This experiment only yields an approximate estimate of the binding affinity since the protein concentration required for sufficient sensitivity was above the Kd for ligand binding. Nevertheless, our data support a binding affinity in the low micromolar range (**Supplementary Figure S14**).

To validate the design models, we solved the structures of two designs, A1E and D2P, by NMR (**Figure 4**). Both designs – A1E in the apo and holo (with progesterone) states, and D2P in the holo state – were in good agreement with the AF2 models. The alpha carbon RMSDs of non-loop regions between the NMR structures and the AF2 models were 1.57 Å for A1E_apo (**Figure 4A**, right) and 1.46 Å for D2P with progesterone (**Figure 4D**). The A1E apo structure was predicted better by the AF2 model than the design model (**Supplementary Figure S15A**). The major difference between the NMR structure to the design model was in helix 3, which moved slightly into the (empty) ligand binding pocket. Interestingly, AF2, which cannot consider a ligand in the prediction, at least partially captured this helix move seen in the NMR structure (**Supplementary Figure S15A**). While we could not determine the ligand pose in A1E_holo due to a lack of NOEs to the ligand (ligand resonances were invisible due to intermediate exchange, also resulting in difficulties determining the conformation of helix 3, **Figure 4B, Supplementary Figure S15B**), plotting the protein combined ^1^H and ^15^N chemical shift differences between the apo and holo states onto the design model agreed with the designed binding pocket (**Figure 4C**). Lastly, the structure of D2P in the presence of progesterone was in good agreement with both the AF2 (**Figure 4D**) and the design models (**Supplementary Figure S15C**), with the main difference occurring in helix 2 (but helix 2 was not designed to contact the ligand binding pocket).

**Figure 4:**
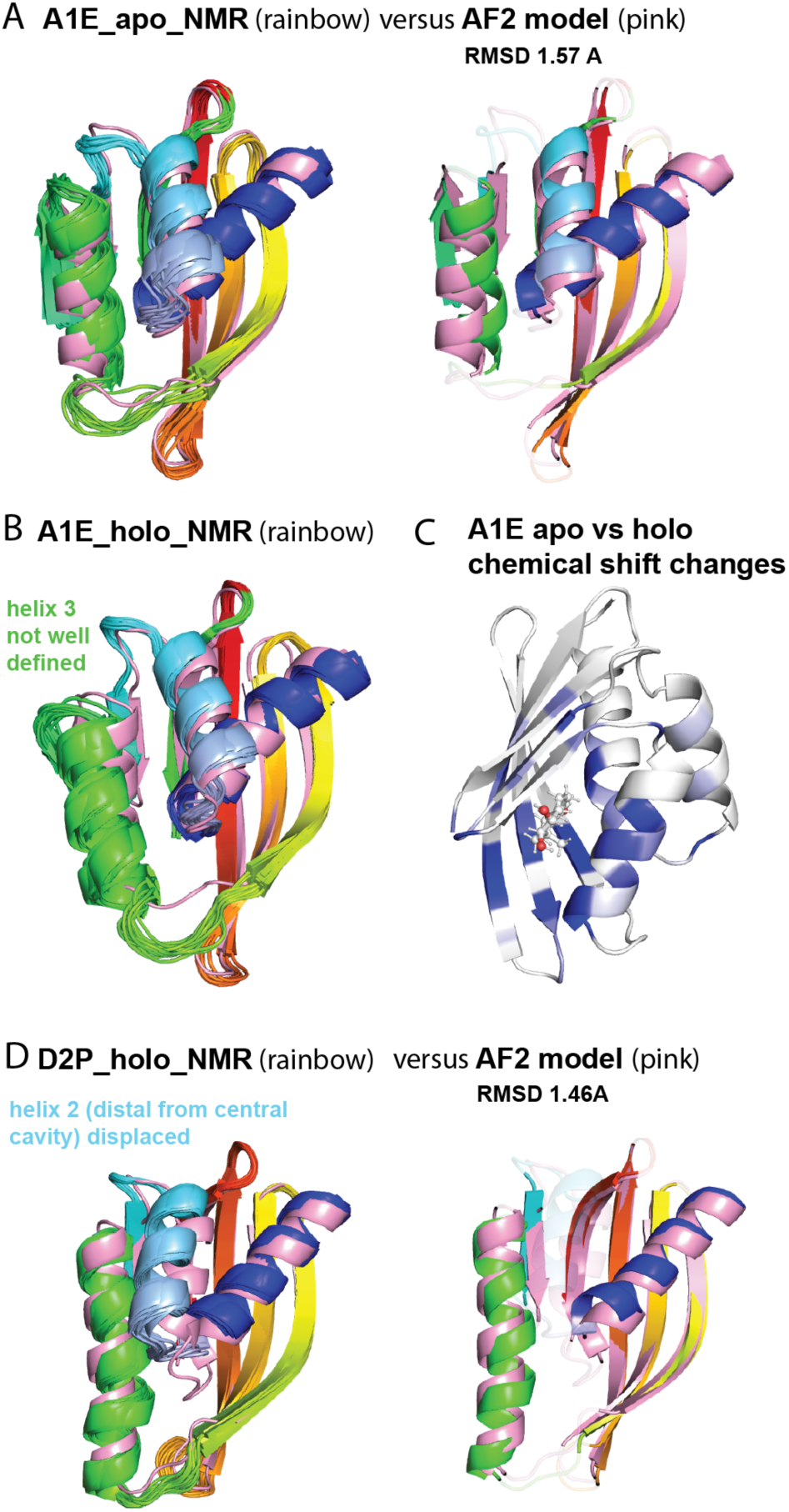
Structural characterization of designed binders. **A**) NMR structure (left, rainbow) for design A1E in the apo state overlaid with the AF2 model (pink) showing good agreement. The alignment with the ensemble average is also shown (right), with non-loop regions used in the RMSD calculation colored. **B**) NMR model of design A1E (rainbow) with progesterone added, overlayed with AF2 model (pink). Helix 3 (green) is not well defined by the NMR data. **C**) NMR chemical shift perturbations plotted on the design model are consistent with the ligand occupying the binding pocket. Darker blue regions indicate larger chemical shift changes upon ligand addition. **D**) NMR model (left, rainbow) for design D2P in the holo state overlaid with the AF2 model (pink). The alignment with the ensemble average is also shown (right), with colored regions (secondary structure elements defining the ligand binding cavity, excluding the distal helix 2) used in the RMSD calculation.

To further validate the designed binding mode, we made individual point mutations for each of these designs targeting polar motif residues defined at the outset of the design workflow to hydrogen bond with the target ligand. We mutated these polar residues to non-polar analogs: N43V, S45V, Y85F for A1E and Y14F, T98V for D2P. All five mutants were monomeric and folded (**Supplementary Figures S16**) and showed a notable decrease in the number and magnitude of ligand-induced chemical shifts in ^15^N HSQC spectra under the conditions tested (**Supplementary Figures S17, S18**), as expected. These results suggest that the designed polar motif residues are critical for binding, supporting the accuracy of the binding mode in our design models.

Finally, we compared the *de novo* designed binding modes to those in human estrogen receptor with estriol and progesterone bound. We find that in all three cases (A1E, A1P, D2P), the designed binders utilize entirely distinct hydrogen bonding and packing arrangements when compared with their naturally occurring counterpart (**Supplementary Figure S19**). In general, the designed binders also have higher protein-ligand interaction density (**Supplementary Figure S19**), supporting the novelty of our design solutions.

## DISCUSSION

The design protocol established in this work synthesizes previously isolated advancements in the generation of protein-small molecule interaction motifs (BSFF) and design of protein scaffolds (LUCS) towards a generalizable workflow for the production of small molecule binding proteins. Building on these tools, we show that using a statistical analysis of BSFF derived interaction modes might help to overcome shortcomings in physical protein design energy functions (such as capturing the geometry of pi stacking interactions), and that employing machine learning methods for sequence design and structure prediction can improve the stability and reliability of LUCS scaffolds. Further, we introduce a strategy for refining the hydrogen bond networks within small molecule binding sites, one of the most difficult features to design well and consequently a major cause of design failure. We show that our refinements increase *in silico* success rates up to ∼7-fold, with design scores across a wide range of commonly used metrics more closely resembling those of naturally occurring small molecule binding proteins.

Finally, we demonstrate the practical utility of our design protocol, applying it towards the production of binders for the steroids estriol and progesterone. In both cases, we generated functional binders with a high experimental success rate, sufficient to eliminate the need for high throughput screening. Our designs are extremely thermally stable, and experimentally determined structures for two designs show good agreement with our design models. While the affinities of our designs for their target molecules could be improved, the efficiency with which we obtain them suggests an immediate use case as a means to obtain starting points for optimization without the need for screening thousands of designs. Conversely, using our design workflow as input to higher throughput methods could drastically increase the number of designs with measurable binding affinity, which would generate informative datasets to improve design methods.

While our methods integrate both machine learning and physics-based tools to capitalize on the relative advantages of each (design stability for the former, explicit functional interactions for the latter), the comparative analysis we provide between diffusion generated designs and our own suggests a potential tradeoff between protein-ligand hydrogen bonding and shape complementarity. Further analysis of the relationship between these features to better understand the source of this tradeoff could inform future efforts to improve design quality.

The results of this work suggest the potential of our workflow for tackling numerous design goals. The general strategy relies on combinatorial sampling from two libraries (binding motifs and protein scaffolds), either of which can be expanded to suit a particular objective. For example, while the NTF2 topology which we used as the basis for our scaffold library frequently binds to hydrophobic molecules (*29*), LUCS could be applied to other protein topologies with a greater proclivity for different classes of molecules, e.g. the GAF domain, which commonly binds to highly polar targets (*30*).

Conversely, while we employed motif libraries on the order of 10^2^, should one need to design a binding site into a particular protein, e.g. to maintain a distinct interface or dynamic behavior, the motif library could be expanded to >10^6^ without compromising the quality of constituent interactions. This adaptability is a key strength of our workflow, and constitutes a significant step towards a general solution for the *de novo* design of small molecule binding proteins.

## METHODS

### Motif library generation

Mol files for the target molecules were downloaded from Pubchem (*31*). The Avogadro molecular editing software (*32*) was used to adjust the hydrogens of the molecules for pH 7.4 and optimize geometries according to the MMFF94 forcefield (*33*). A python script included with Rosetta, ‘molfile_to_params.py,’ was used to generate Rosetta params files for target molecules. A custom Python script, ‘Fragment_Search.py’ was then used to identify molecules in the PDB containing user defined fragments as substructures, with the fragments being provided as lists of corresponding atom names in an input text file.

All residues within 4 or 3.5 angstroms of the substructures for non-polar and polar residues, respectively, were then extracted from relevant PDB entries and transformed relative to the target molecule. Redundant entries were discarded based on sequence identity of extracted residues, with a cutoff of 70%. Any residues with missing atoms were also discarded. The residues in these filtered interaction pools were then spatially clustered, with each cluster containing residues of a single amino acid type with mutual RMSD within 0.5 Å (**Figure 1D**, **Supplementary Figure S1**).

Motifs were described by a Rosetta match cst file, which allows for definition of six degrees of freedom for each desired interaction. The motifs were generated by first specifying idealized hydrogen bonding interactions for each polar atom in the target molecule for which a single hydrogen bond was sufficient (e.g. an isolated hydroxyl oxygen), with the ideal value and tolerance for each parameter obtained from a statistical analysis of protein-ligand interaction preferences in the PDB. The full set of unique residue-polar atom combinations was enumerated, with restrictions on redundancy (no more than 2 of the same residue type per motif), and limiting the number of multi-polar atom residues (arginine, asparagine, glutamine) per motif to 1.

Randomly selected non-polar packing interactions were then added to a random subset of these hydrogen bonding interactions, using only those non-polar interaction modes that were overrepresented in the clustering analysis (top 10th percentile of cluster population) and ensuring that the selected interactions did not clash with the target molecule. The interaction geometries were derived from a representative member of the source cluster and were not allowed to deviate from the observed values in the representative member beyond 0.25 Å distance for the interacting atoms and 15 degrees for all relevant angles (**Supplementary Figure S2**) so as to preserve the desired interaction mode. Rosetta was used to score these interactions and ensure no clashes with the ligand or any of the other existing motif residues. A subset of pairwise energy terms was employed to evaluate the motifs; fa_atr, fa_rep, hbond_sc, hbond_bb_sc, fa_elec; using the same weights as in the standard Rosetta energy function (1, 0.55, 1, 1, 1, respectively). This same scoring function was used for the plot of score distributions of tryptophan-benzene clusters in **Figure 1D**.

For certain polar fragments with strong observed preferences for higher order hydrogen bonding, such as carboxylate groups, neighboring hydroxyls, guanidinium nitrogens, etc., explicit interactions were added to the motifs in a manner analogous to that of the non-polar packing interactions, with the exception that the pool of clusters from which interactions were chosen was limited to those which simultaneously formed two hydrogen bonds with the ligand fragment. For example, two nitrogens from an arginine each hydrogen bonding with one of the oxygen atoms in a ligand carboxylate group.

This process resulted in 230 motifs for estriol and 399 motifs for progesterone, with all consisting of 3 hydrogen bonds and 1 non-polar packing interaction. The desired number of motifs generated is user defined for each case, with typically hundreds of possible interactions for a given fragment, and N Choose M possible motifs without redundant interactions for an M-residue motif, where N is the size of the interaction pool to select from.

For testing the generalizability of the protocol, 50 4-residue motifs each were also assembled for caffeic acid (DHC), 3-cyano-7-hydroxy-coumarin (3QV), and DFHBI (38E) according to the same strategy described above (prioritization of hydrogen bond interactions with ligand polar atoms). For the examples shown in **Figure 1E**, these motifs were grafted into the 1,816 de novo NTF2 scaffolds described below, and only well-buried matches were taken (as described below, motif grafting).

### Protein scaffold library generation

We began with 1,709 LUCS designs with two reshaped helices on an input *de novo* designed NTF2 protein (PDB 5TPJ) generated as described previously (*33*).

Natural NTF2 folds contain a central cavity to bind a variety of ligands. These designs were filtered to ensure a lack of buried unsatisfied hydrogen bonds, high shape complementarity in the reshaped region, and high local sequence-structure compatibility (Rosetta Fragment Quality ≤ 1.0 Å, average RMSD between reshaped region and sequence similar fragments from crystal structures). For each generated backbone, we produced 400 sequences using protein MPNN, and predicted the structure for the top 10 scoring sequences using ColabFold, retaining those designs with an average pLDDT greater than 88.0 and a Ca RMSD between the design model and predicted structure of less than 1 Å. In cases where more than 1 of the 5 ColabFold models for a given input sequence met these criteria, we retained only the one with the highest confidence. We then applied the Rosetta FastRelax algorithm to these filtered ColabFold models and submitted them again for sequence redesign using ProteinMPNN, repeating this entire process over a total of 3 rounds. In the final round, in addition to the ColabFold metrics, we also filtered the designs by Rosetta metrics including quality of atomic packing (Packstat ≥ 0.65), hydrophobic SASA ≤ 58% of total SASA, and less than 4 buried unsatisfied hydrogen bonds. This protocol resulted in 1,816 scaffolds with an average RMSD in the reshaped regions of 6.5 Å with the input 5TPJ (**Supplementary Figure S20**).

### Motif grafting into protein scaffolds

We used the Rosetta Match application to scaffold our motifs into backbones from our NTF2 scaffold library, employing expanded rotamer sampling (options - ex1, - ex2, -extrachi 5). The ideal values for the 6 degrees of freedom used to specify a desired interaction were either taken directly from the observed values in a representative of the clustered interaction mode being recapitulated (see section ‘Motif library generation’), or in the case of simple hydrogen bonds, were derived from a statistical analysis of hydrogen bonding geometries across all BSFF derived protein-ligand hydrogen bonds. The tolerances for these values were narrowly defined so as to preserve the desired interaction geometries (**Supplementary Figure S2**). The torsion, angle, and distance constraint sampling numbers were 5, 3, and 1 respectively for simple hydrogen bonds, and 2, 2, 0 for other interaction types. These values specify the number of points to sample between the ideal value (which is always sampled by default) and the defined tolerance, such that the total number of sampled values is 2N+1, where N is the constraint sampling number. For example, for an ideal distance of 1.9 Å with a tolerance of 0.5 and a sampling number of 2, the distances sampled would be 1.4, 1.65, 1.9, 2.15, and 2.4 Å.

Raw matches were filtered according to ligand burial using the Rosetta dsasa metric, which describes the fraction of the ligand surface area that is buried upon binding, using a value of 0.7 as the cutoff to identify well buried ligands.

### Binding site sequence design

Rosetta resfiles, which specify the allowed residues at each position during sequence design, were created for the filtered matches, allowing design to all non-cysteine residues for any amino acid with heavy atoms within 5 Å of the ligand.

Sequence design was then carried out for each match using the Rosetta Enzyme Design application (*34*), employing the beta_nov16 score function (*35*) with protein-ligand interactions up-weighted by 2.5 relative to protein-protein interactions over 3 cycles of design and minimization. Designs were then filtered, excluding those with high ligand hydrophobic SASA (>40 Å^2^), oversaturated ligand polar atoms, and poor per residue energies (≥-2.0 Rosetta energy units (REU)) for residues in the binding site.

### Binding site hydrogen bond refinement

For designs passing initial metrics, a custom python script was used to refine polar residues within the binding site, ‘hbrefine.py.’ This script first identified 2 features which often lead to failed designs, extraneous polar residues in the binding site (i.e. not hydrogen bonding with ligand or protein) and unsatisfied ligand polar atoms. For the former, identified residues were mutated to non-polar residues, excluding alanine (leucine, isoleucine, valine, phenylalanine), repacked, and then accepted if the Rosetta energy score was equal to or lower than that of the original residue (if multiple such mutations were identified, the lowest scoring mutation was kept). For unsatisfied ligand polar atoms, all adjacent residues were mutated to serine, threonine, lysine, tryptophan, and tyrosine, and then repacked. These mutations were accepted if they formed a new hydrogen bond with the unsatisfied ligand polar atom and improved or maintained the Rosetta energy score of the design.

These refined designs were then filtered according to the previously described metrics (ligand hydrophobic SASA, oversaturated polar atoms, per residue binding energy), along with additional requirements of having at least 3 hydrogen bonds with the ligand, having no unsatisfied buried polar atoms in the binding site, high protein-ligand shape complementarity (≥ 0.6), and high binding site pre-organization (average binding site rotamer Boltzmann probability ≥ 0.3).

### Stability oriented sequence design

Next, ProteinMPNN was used to redesign all residues outside of the binding site (all residues which were not designable in ‘Binding site sequence design’ above) in order to improve the sequence-structure compatibility of the designs and stabilize the binding site, which at this point is substantially altered relative to the initial scaffold sequence. A sampling temperature of 0.15 was employed, and 300 sequences were produced for each input protein. The Rosetta FastDesign algorithm was then applied to these same input proteins, allowing design to only those amino acids observed within the ProteinMPNN sequence profiles at each position. Up to ten designs were made for each input, and then filtered according to the previously described binding site-oriented metrics (see ‘Binding site hydrogen bond refinement’ section above), as well as additional stability focused metrics, including high packing density (Rosetta PackStat ≥ 0.6), low cavity volume in the holo state (≤120 Å^3^), and low exposed hydrophobic surface area (≤1200 Å^2^).

### Final filtering of designs

Designs passing all criteria were submitted to ColabFold for structure prediction, and then filtered to those with an average pLDDT value greater than or equal to 88.0 and an alpha carbon RMSD between input design model and predicted structure of less than 1.5 Å.

Finally, the resultant designs were analyzed visually to check for large hydrophobic surface patches. In certain cases, point mutations were introduced using standalone FoldIt (*36*) to address such issues, and the resultant mutants re-scored by Rosetta to ensure retention of all previously described quality criteria. Four such designs were amongst those ordered for experimental screening, A1P, B1P, C1P, and D1P, which contained mutations W93T, W80K, W113S, and Y99K, respectively.

### Production runs

A total of 26 designs were ordered for experimental screening, 13 for each of estriol and progesterone. Designs targeting progesterone were obtained according to precisely the workflow described in the previous sections. Designs targeting estriol, however, were generated in conjunction with the development of this finalized workflow. Consequently, the methodology used to obtain these designs differed in several ways. Firstly, an earlier iteration of the refined LUCS NTF2 scaffold library was used for matching, containing only 515 structures, rather than 1,816. Second, an initial production run with the resultant matches occurred prior to the development of our HBRefine protocol, with this producing 9 out of the 13 ordered design sequences (A1-B2). After HBrefine development, a subset of the original estriol matches was subject to the finalized workflow (with the inclusion of HBRefine), producing 4 additional sequences for experimental testing (C2-G2).

### Derivation of *in silico* filtering criteria

The selection of quality criteria used for *in silico* filtering of designs was informed by analysis of existing monomeric small molecule binding proteins in the BindingMOAD database (*36*), which had both an experimentally determined binding affinity and a refined, high quality structure present in the PDBredo database (*37*). The final dataset included 366 structures with 312 unique ligands (available in **Supplementary Materials**). These structures were relaxed into the Rosetta energy function using the standard FastRelax protocol with the beta_nov16 all-atom scoring function, along with the following flags:

-extra_res_fa LIGAND_PARAMS.params -ex1 -ex2 -extrachi_cutoff 0 -nstruct 1 - in:file:fullatom -relax:fast -relax:constrain_relax_to_start_coords -relax:coord_cst_stdev.5

and then evaluated according to the metrics listed in **Supplementary Table S1**. While no significant correlations were found between binding affinity and any of the individual metrics analyzed, score distributions for these known binders showed near universally strong scores across all binding site features, suggesting that, while no one metric may be predictive of binding, strong scores across many metrics simultaneously are. We thus employed a range of diverse filtering criteria in our design workflow, requiring simultaneous satisfaction of all. The cutoff values we employed were informed by the average normalized values across the known binders (see **Supplementary Figure S4**), either using the average value directly or, in certain cases, relaxing the threshold slightly while ensuring it to still be within the range of observed values for the MOAD binders (e.g. per-residue exposed hydrophobic surface area cutoff was 10.5, compared to an average value of 8.5 across MOAD binders).

### Design with RosettaFold Diffusion all-atom

For the RFdiffusion All Atom design simulations, progesterone and estriol models were first idealized in Avogadro by adding hydrogens according to a pH of 7.4 and minimizing energy using the MM94 forcefield, as described above for designs generated using our method. RFdiffusion All-Atom was run with parameters identical to the Krishna et al. (*37*) digoxigenin binder design pipeline to generate 6,000 starting backbones.

Backbones were filtered for ligand SASA ≤ 100, radius of gyration ≤ 13, and loop ratio < 0.4, resulting in 3,968 backbones for progesterone and 3,585 for estriol.

Three rounds of LigandMPNN and Rosetta Fastrelax were then performed to iteratively optimize the sequence-backbone combinations. In round one, 8 sequences were designed per backbone resulting in 31,744 sequences for progesterone and 28,680 for estriol, and in subsequent rounds one sequence was generated per input. Between rounds, the LigandMPNN packed model was relaxed with Rosetta cartesian FastRelax. Designs were then analyzed using AlphaFold2 in single sequence mode and Rosetta for self-consistency and previously described *in silico* quality metrics.

### Design test sets for comparison of *in silico* success rates

For comparisons of *in silico* success rates, both within our own design workflow (**Figure 1A-C**) and with RFDaa generated designs (**Figure 2**), we used test sets of estriol and progesterone binder designs containing 8,500 well-buried matches. The estriol matches included 78 unique motifs and 1,612 unique scaffolds. The progesterone matches included 17 unique motifs and 531 unique scaffolds. The matches were subject to our full design workflow, and the outputs from each step (binding site design, hydrogen-bond refinement, and ProteinMPNN scaffold stabilization) were used for the determination of *in silico* success rate improvements from refinement (**Figure 1G**, **Supplementary Table S7**). The final sets of unfiltered designs after the ProteinMPNN design step were used for comparison with estriol and progesterone binders from RFDaa (**Figure 2A-D, Supplementary Table S8**).

### Matching with motifs and scaffolds available in PDB

NTF2 scaffolds with available crystal structures were identified by querying the Protein Data Bank using the InterPro identifier for the NTF2-like domain superfamily (IPR032710) and filtering the results to only single domain entries (number of polymer instances = 1) with sequence length ≤ 150 residues. The results were grouped by sequence identity ≤ 70%, and the structure in each group with the highest resolution was taken as representative.

Motifs were constructed by first identifying all structures in the PDB with estriol or progesterone bound. These were then processed to extract only one chain with the relevant molecule bound. Motifs were assembled by first identifying all protein-ligand hydrogen bonds and prioritizing their incorporation. Additional interactions were added to the desired total number of motif residues by taking the residues with the lowest pairwise protein-ligand Rosetta energy scores.

An initial matching run was performed using motif parameters mirroring those of the estriol and progesterone design runs with CLAIRE (4 residue motifs, interactions constrained within 0.25 Å distances, angles within 15°) but did not produce any matches in either case. A subsequent matching run was performed using only 3 motif residues, and relaxing the interaction distance constraint to 0.5 Å. This produced 10 well buried matches for estriol and 210 for progesterone. The Rosetta EnzymeDesign application was used to produce 10 designs for each of these matches, with none passing our first round of binding site evaluation criteria.

### Protein Expression

pET-28a+ plasmids including the designed sequences were ordered from Twist Bioscience, with the sequence for the designs inserted between the Ndel and Xhol restriction sites, including an N-terminal polyhistidine tag.

*E. Coli* BL21 (DE3) cells were transformed with these plasmids and cultured overnight at 37 °C. Colonies were then inoculated into 5 mL of LB media and cultured overnight in a shaker at 37 °C, 225 RPM. Cells were pelleted at 18,000 x g for 10 minutes and then resuspended into 1.5 mL of PBS buffer at pH 7.4 (135 mM NaCl, 2.6 mM KCl, 10 mM Na_3_PO_4_, 1.8 mM KH_2_PO_4_). The cells were lysed using B-PER solution from Thermo Fisher and the resultant lysate tested for expression and solubility via Coomassie stain SDS-PAGE.

Bacterial cultures expressing soluble designs were then inoculated into 1L of LB media and grown in a shaker at 37 °C, 225 RPM, until reaching an optical density of 0.6. 50 µg/mL of isopropyl β-D-1-thiogalactopyranoside (IPTG) was then added to these cultures, and they were placed back in shakers for four hours. Cells were again pelleted and lysed, and soluble and insoluble fractions were separated by centrifugation at 20,000 x g for 20 minutes. Soluble fractions were mixed with Ni-resin beads to purify the designed proteins by Immobilized Metal Affinity Chromatography (IMAC). The beads were washed with 5 x 1 mL wash buffer, followed by elution with 3 x 1 mL of elution buffer.

### Analytical Size Exclusion Chromatography

Purified protein samples were analyzed using a Superdex® 75 10/300 GL size exclusion column with PBS buffer. The relationship between elution time and log molecular weight was determined via application of a linear regression model to the elution times of the proteins from the BioRad Gel Filtration Standard, and the oligomeric state and molecular weight of the designed proteins was determined by comparison of their respective elution times to the fit of the standard.

### Circular Dichroism Spectroscopy

Circular dichroism (CD) data were collected using a Jasco J-710 spectrometer. The proteins were diluted to a concentration of ∼15 µM in PBS buffer, and spectra were measured at 25 °C in a 1 mm cuvette, scanning between wavelengths 200-280 nm.

Melting curves were then collected at 220 nm, scanning temperature between 25 and 95 °C at a rate of 1 °C per minute.

### 15N HSQC Spectra

Purified proteins were prepared in PBS to a final concentration between 100-300 µM, depending on yield. D_2_O was added such that the final volume had 5% D_2_O and 95% H_2_O. For spectra taken with ligand, the ligands were first solubilized in organic solvent (either dimethylformamide or dimethyl sulfoxide), then diluted with PBS to a composition of 50% solvent, co-solvent by volume. The appropriate volume was added from this solution to the experimental sample, such that the final sample contained 5% organic co-solvent by volume. The estriol concentrations of the samples used were generally between 400-600 µM, while the progesterone concentrations were between 500 µM-1.06 mM (see Figure captions for relevant conditions).

### Protein expression for NMR structure determination

^15^N and ^13^C labeled proteins were expressed by growing *E. coli* in M9 minimal medium that included ^13^C-glucose and ^15^NH_4_Cl (6g/L Na_2_HPO_4_, 3g/L KH_2_PO_4_, 0.5g/L NaCl, 0.5g/L ^15^NH_4_Cl, 50mg/L EDTA, 8.3mg/L FeCl_3_ x 6 H_2_O, 0.84mg/L ZnCl_2_, 0.13mg/L CuCl_2_ x 2 H_2_O, 0.1mg/L CoCl_2_ x 6 H_2_O, 0.1mg/L H_3_BO_3_, 0.016mg/L MnCl_2_ x 6 H_2_O, 0.2% (w/v) ^13^C-glucose, 1mM MgSO_4_, 0.3mM CaCl_2_, 1mg/L Biotin, 1mg/L Thiamine). Single bacterial colonies were first inoculated into 5 mL seed culture and grown at 37 °C overnight. The seed culture was inoculated into 1L ^15^N ^13^C labeled M9 minimal medium and grown at 37 °C until OD600 reached 0.5-0.7. Then 1 mL 1M IPTG was added to induce protein expression at 30 °C overnight. The expressed proteins were purified following the Ni-resin pull down protocol described in the protein purification section. For design D2P, the N-terminal His tag was cleaved by overnight digestion with thrombin at 25 °C and further purified using a HiLoad® 16/600 Superdex® 75 pg size exclusion column from GE with 50mM phosphate buffer at pH 7.0. The monomeric fractions were collected and concentrated to 0.5-1 mM in PBS at pH 7.4 for NMR experiments. For design A1E, the His tag was not cleaved for structure determination.

### Structure determination by NMR

The final protein concentrations were approximately 500 μM. NMR spectra were measured at 298.1K. Two dimensional (2D) ^1^H, ^15^N-HSQC (pulse program: fhsqcf3gpph), 2D ^1^H,^13^C-HSQC (pulse program: hsqcetgpsisp2), XXms 3D CCH-TOCSY (pulse program: XXX) and XXms 3D simultaneous ^13^C/^15^N-NOESY-HSQC (pulse program: noesyhsqcgpsismsp3d) spectra were measured using a Bruker NEO 800 MHz spectrometer with a 5mm TCI H&F-C/N-D CryoProbe. 2D TROSY (pulse program: trosyargpphwg), 3D ^1^H-^13^C NOESY-TROSY (pulse program: noesytrosyargpphwg), 3D CACB(CO)NH (pulse program: hncocacbgpwg3d), 3D CACBNH (pulse program: hncacbgpwg3d), 3D Hcc(co)NH (pulse program: hccconhgpwg3d2), 3D Cc(co)NH (pulse program: hccconhgpwg3d3), 2D HBCBCGCDHDGP (pulse program: hbcbcgcdhdgp), and 2D HBCBCGCDCEHEGP (pulse program: hbcbcgcdcehegp) were collected on a Bruker Avance 600 MHz spectrometer with an Inverse 5mm H-C/N-D cryoprobe. Spectra were processed in TopSpin 3.6.3.

Automated peak picking, resonance assignment, and structure calculations were performed by ARTINA (*38*), and dihedral angle restraints were generated by TALOS (*39*). Manual inspection of ARTINA chemical shift assignments was done in CCPNMR v.3.1.0 (*40*) and corrections were made as necessary. The manually curated chemical shift list was then included as input for a subsequent structure calculation run in ARTINA. The output candidate structure (an ensemble of n=20 conformers) with the lowest CYANA target function value was refined in XPLOR-NIH-3.7 with the refine.py script included in the distribution (eginput/gb1_rdc/refine.py) (*41*) (*42*). The 20 lowest scoring structures out of 100 were then refined in explicit water with the wrefine.py script (eginput/gb1_rdc/wrefine.py). The ensemble of the refined structures was validated using the PDB validation server.

### NMR titrations

Protein and ligand stocks were prepared as previously described, with the final composition of the ligand stock used in titration experiments containing 500 μM progesterone in sodium phosphate buffer pH 7.4, with 1% DMF and 5% D2O. The initial samples for titration experiments contained 72 μM protein in 1mL total volume sodium phosphate buffer pH 7.4, with 1% DMF and 5% D2O. For subsequent titration points, ligand stock solution was added to the experimental sample at volumes of 10, 20, 30, 60, and 120 μL, producing progesterone concentrations of 4.95, 14.5, 28.3, 53.5, and 96.7 μM, respectively, and diluting the protein concentration to 71.3, 69.9, 67.9, 64.3, and 58.1 μM.

2D ^15^N heteronuclear single quantum coherence (Fast HSQC (*43*)) spectra (TopSpin 4.3.0 pulseprogram: fhsqcf3gpph) were acquired at 25 °C on a Bruker Avance NEO 800 MHz spectrometer equipped with a 5 mm TCI-Cryoprobe with actively shielded Z-axis gradients.

Eight scans were acquired per complex t1 point with 128* and 640* complex points and spectral widths of 2,838 Hz and 12,500 Hz for the ^15^N and ^1^H dimensions, respectively. Spectra were processed using Bruker TopSpin 4.3.0 software. The number of points in the ^15^N dimension was doubled with linear prediction, zero-filled and apodised with a 90 deg shifted squared sine-bell window function (QSIN), before Fourier transform and base-line correction (proton dimension). Peak intensities were extracted using the CCPN package (*15*, *43*).

### Similarity of designed motifs to interactions with ESL and PRG in natural proteins

Structures of human estrogen receptor with estriol and progesterone bound were downloaded from the Protein Data Bank (structure codes 3Q95 and 1A28, respectively). Analysis of binding site properties for comparison with those of our designed binders was performed using the Protein Plus webserver (https://proteins.plus) (*44–47*). The PoseEdit tool was used to produce binding site interaction schematics (*48*), while pocket features were obtained using the Geomine tool (*49*) (*50*, *51*).

## Acknowledgements

We thank the Kortemme group, Lee Schnaider, and Chad Altobelli for helpful discussion.

## Author contributions

C.V.G. and T.K. developed the conceptual approach. C.V.G. developed the computational design methods and performed and analyzed the design simulations. C.V.G. and I.L.A characterized designed proteins experimentally, with contributions and advice from D.G. C.V.G., I.L.A., D.G. and M.J.S.K. collected and processed NMR data. A.B.G. solved the NMR structure of A1E with guidance from M.J.S.K.. M.N.C. performed RFDiffaa-based design and Boltz-2 calculations. S.K. and D.K. contributed to computational design simulations. T.K. provided guidance and resources. C.V.G. and T.K. wrote the manuscript with contributions from the other authors. All authors read and commented on the manuscript.

## Code and data availability

The collection of python3 scripts used for the design and analysis presented in this work is available at https://github.com/cvgalvin/CLAIRE. A readme file therein contains detailed instruction for reproducing the design workflow or extending it to novel targets. Provided in https://github.com/cvgalvin/CLAIRE/tools are the xml scripts for Rosetta protocols implemented in the main code, which include all relevant parameters, along with the commands and parameters used for RosettaFold Diffusion all-atom, ProteinMPNN, ColabFold, and Boltz-2. The NMR structure of A1E will be deposited to the Protein Data Bank, PDB code 11EZ.

## SUPPLEMENTARY FIGURES

**Supplementary Figure S1:**
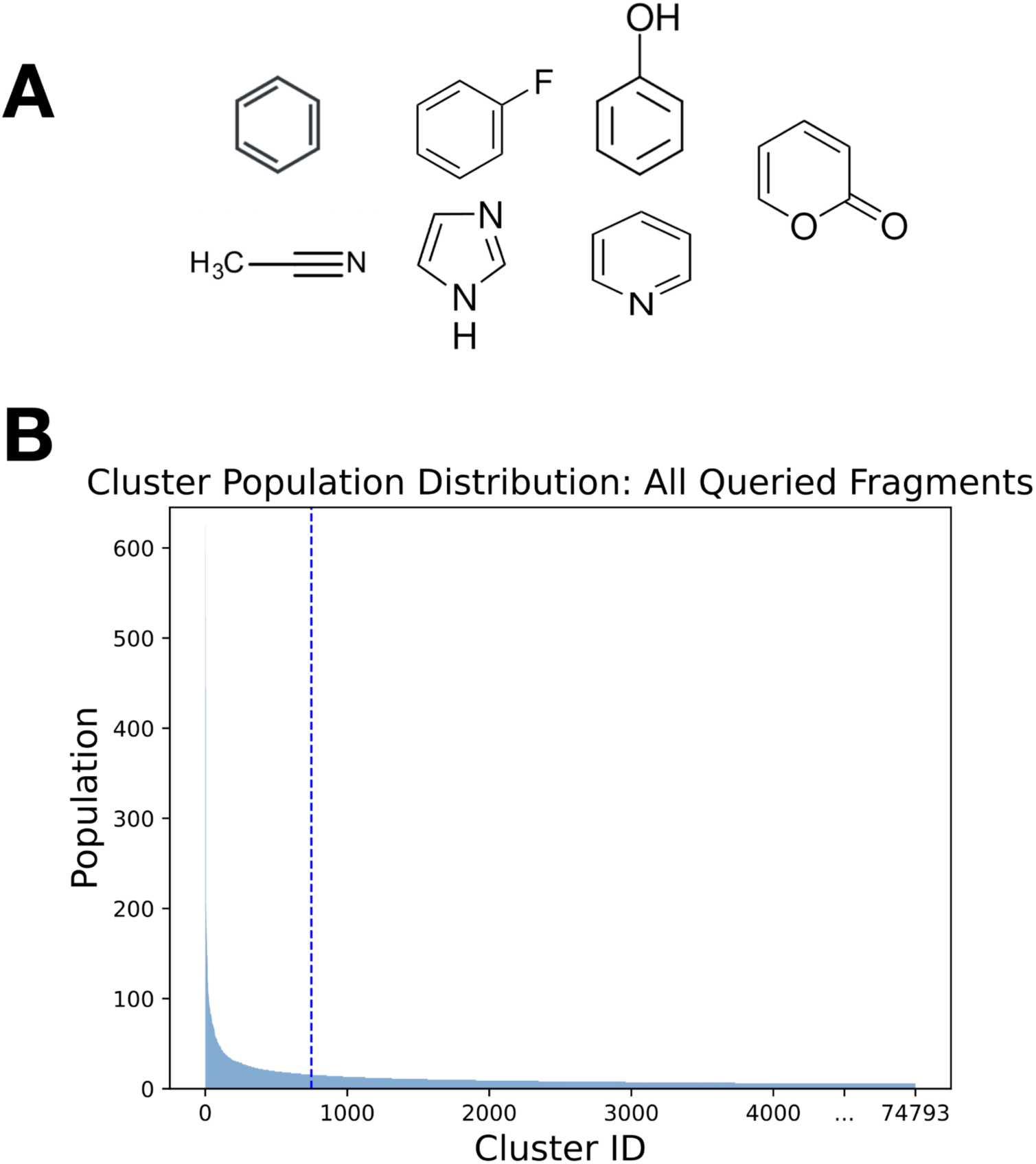
Cluster population distribution across queried ligand substructures. A) Chemical structures of queried ligand fragments. B) Distribution of cluster populations, ordered from most to least highly populated, for ligand chemical substructures shown in panel A. The top 1% most frequently observed interaction modes are indicated by a dashed blue line and account for ∼15% of all observed contacts in this set.

**Supplementary Figure S2:**
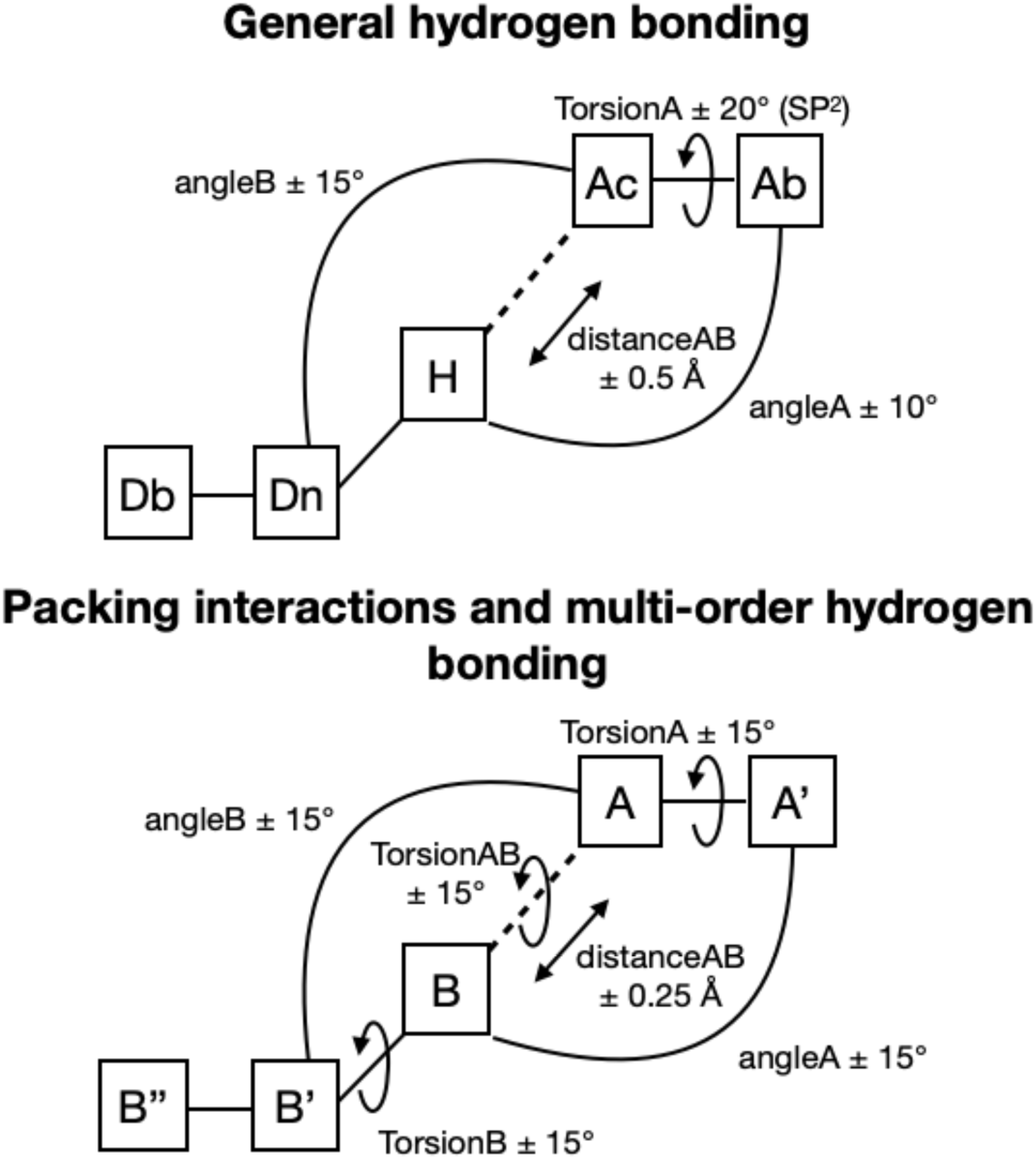
Narrow tolerance windows ensure high fidelity between defined interactions and grafted interaction geometries. The ideal values for each parameter vary with the interaction, but in all cases the geometries are strictly bound to preserve the desired interaction mode. In addition, designs were minimized using the Rosetta force field and FastRelax protocol, as described in Methods, and filtered using the Rosetta hydrogen bond potential that has stringent geometry parameters for distance and angles (*15*, *52*). Top: relevant parameters for hydrogen bonds. Ac: acceptor atom. Ab: atom bound to acceptor. H: hydrogen. Dn: donor atom. Db: atom bound to donor atom. Bottom: General definition of relative orientation of three atoms on the protein (B, B’ and B’’) and ligand (A, B’ and A’’) defined by six degrees of freedom.

**Supplementary Figure S3:**
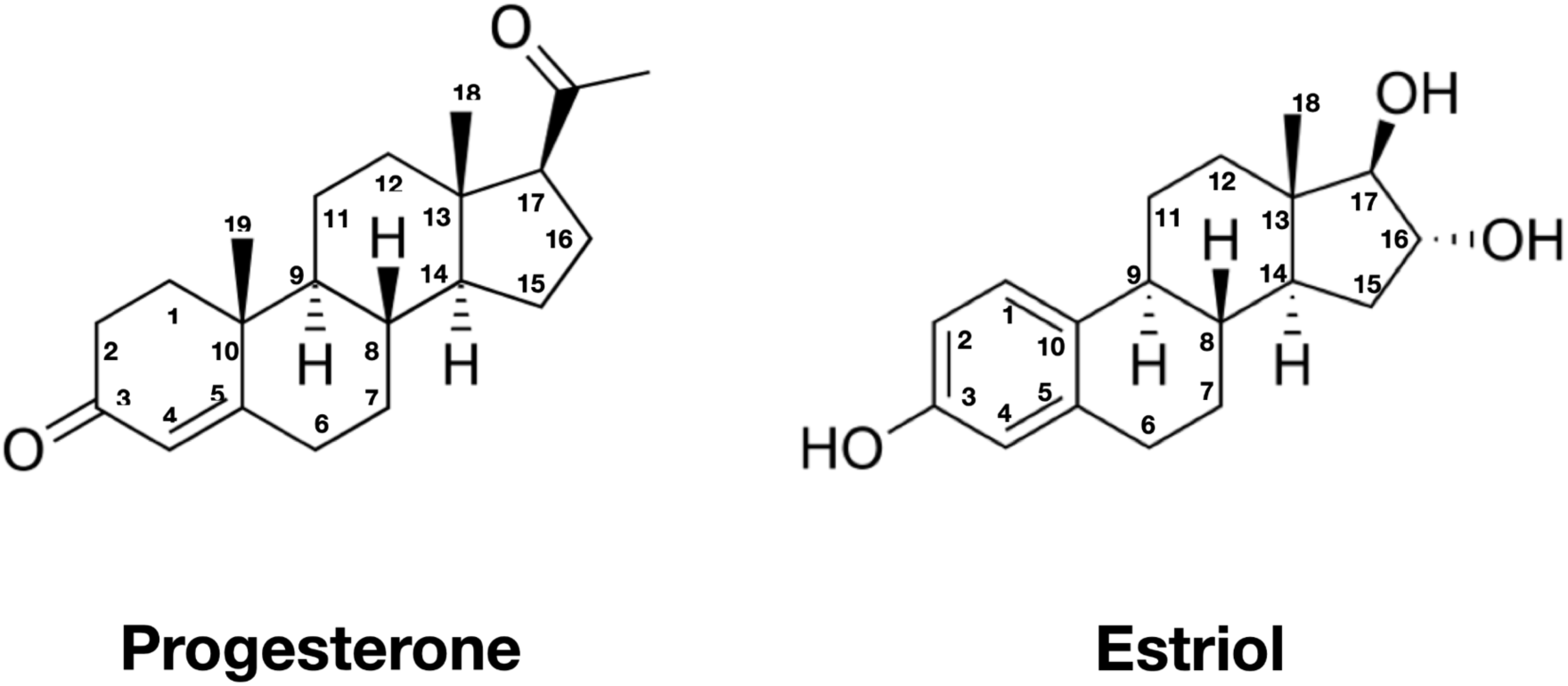
Steroid design targets progesterone and estriol, with atom numberings. The key differences are the substituents at the 3,16, and 17 positions. Progesterone has a carbonyl, no substituent, and an ethyl ketone at each of these positions, respectively, while estriol has 3 hydroxyl groups. Progesterone also possesses an additional methyl substituent at the 10 position.

**Supplementary Figure S4:**
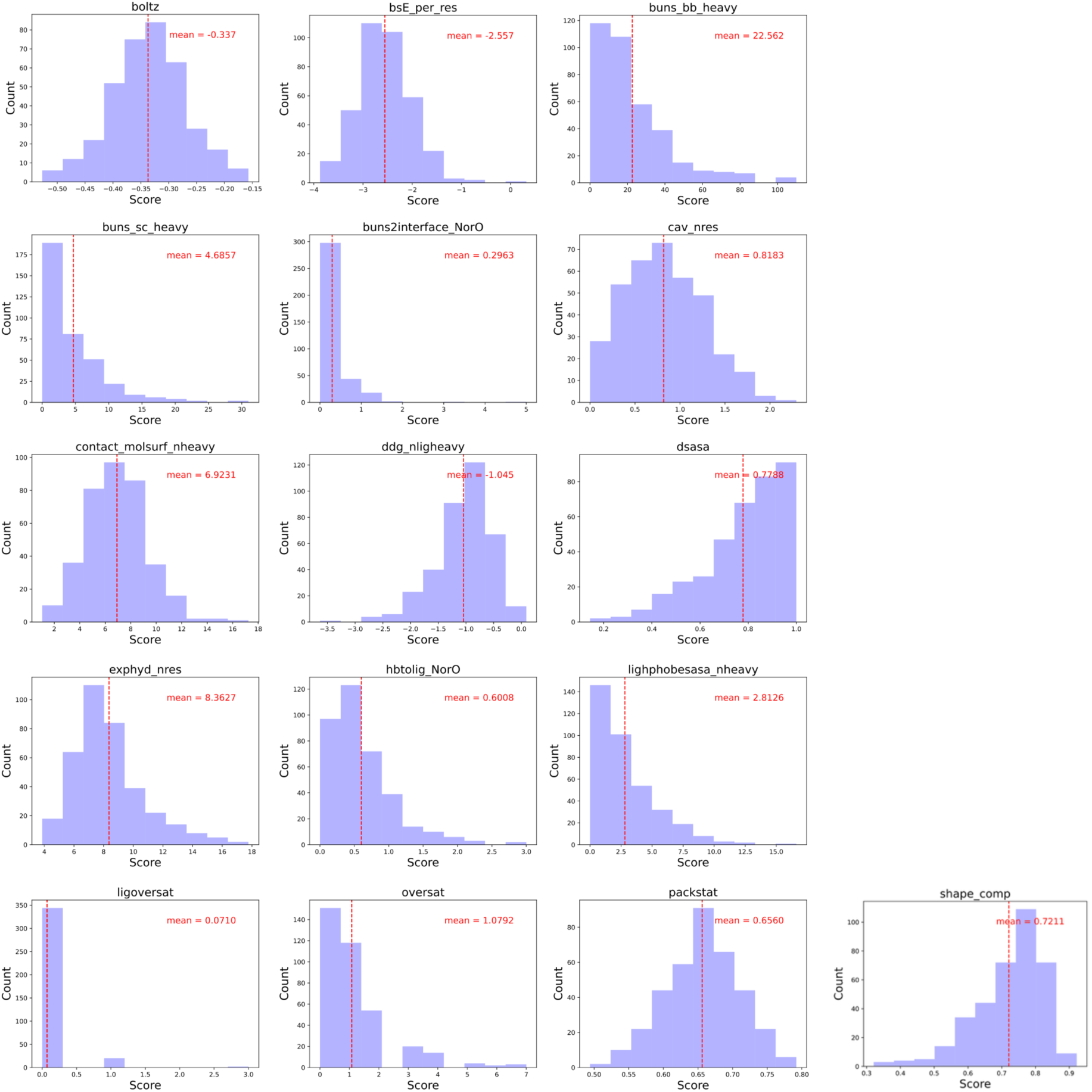
Distributions of scores for structures in the BindingMOAD dataset. The distributions are shown as histograms, with the mean values indicated with a dotted line as well as a text overlay. The metric names correspond to those described in **Supplementary Table S1**, where their meanings can be found. Suffixes on terms indicate normalization factors, e.g. ‘NorO’ indicates the raw score was normalized by the number of nitrogen and oxygen atoms in the ligand, ‘nres’ indicates that the raw score was normalized by the total number of residues in the protein, ’nheavy’ indicates that the raw score was normalized by the total number of ligand heavy atoms.

**Supplementary Figure S5:**
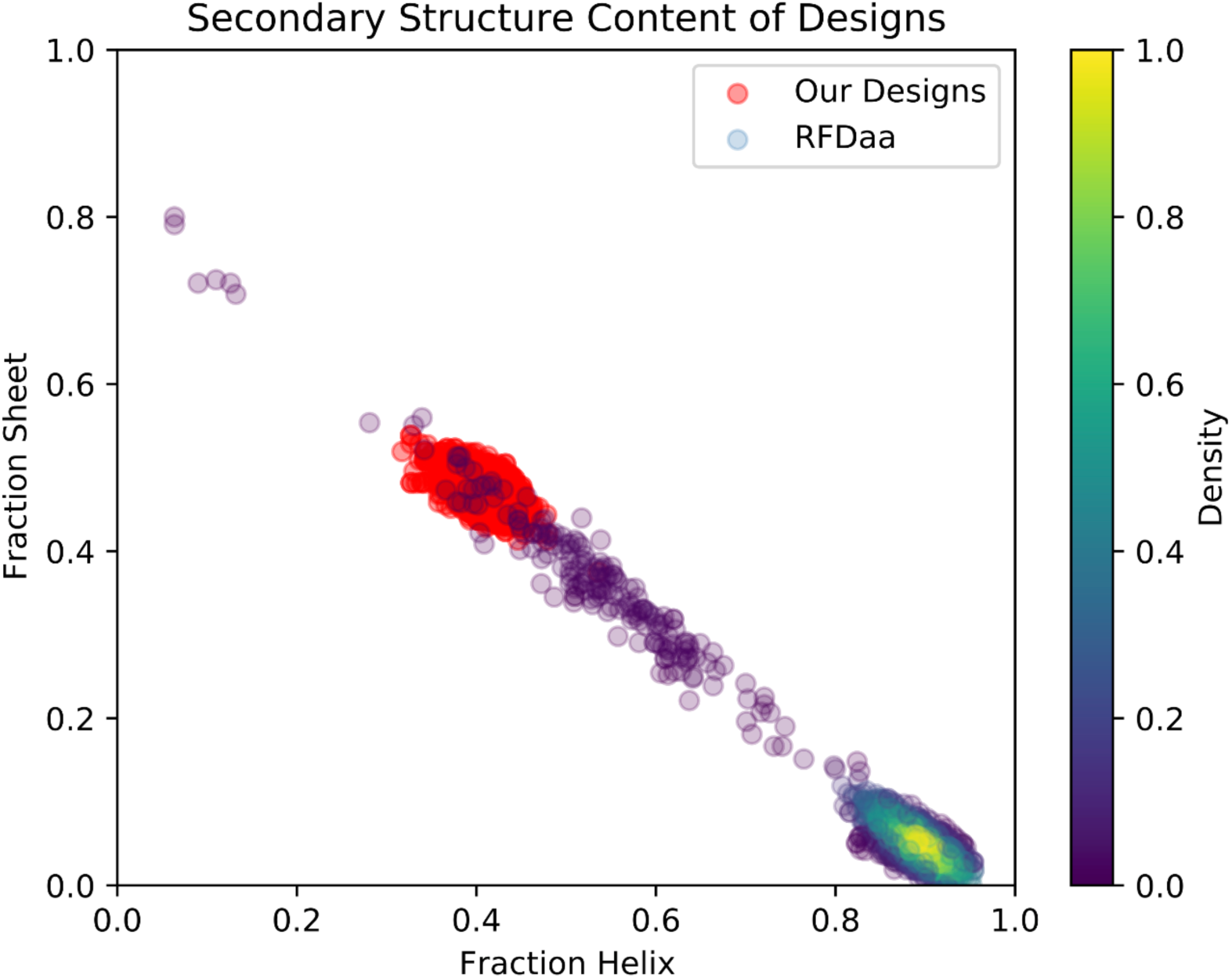
Secondary structure content across our design models and RFDaa generated design models for both estriol and progesterone binders. The distribution for our designs is shown in red, while for RFDaa designs it is colored by point density, showing that they are concentrated mainly in the lower right region of the plot, which represents helical bundles.

**Supplementary Figure S6:**
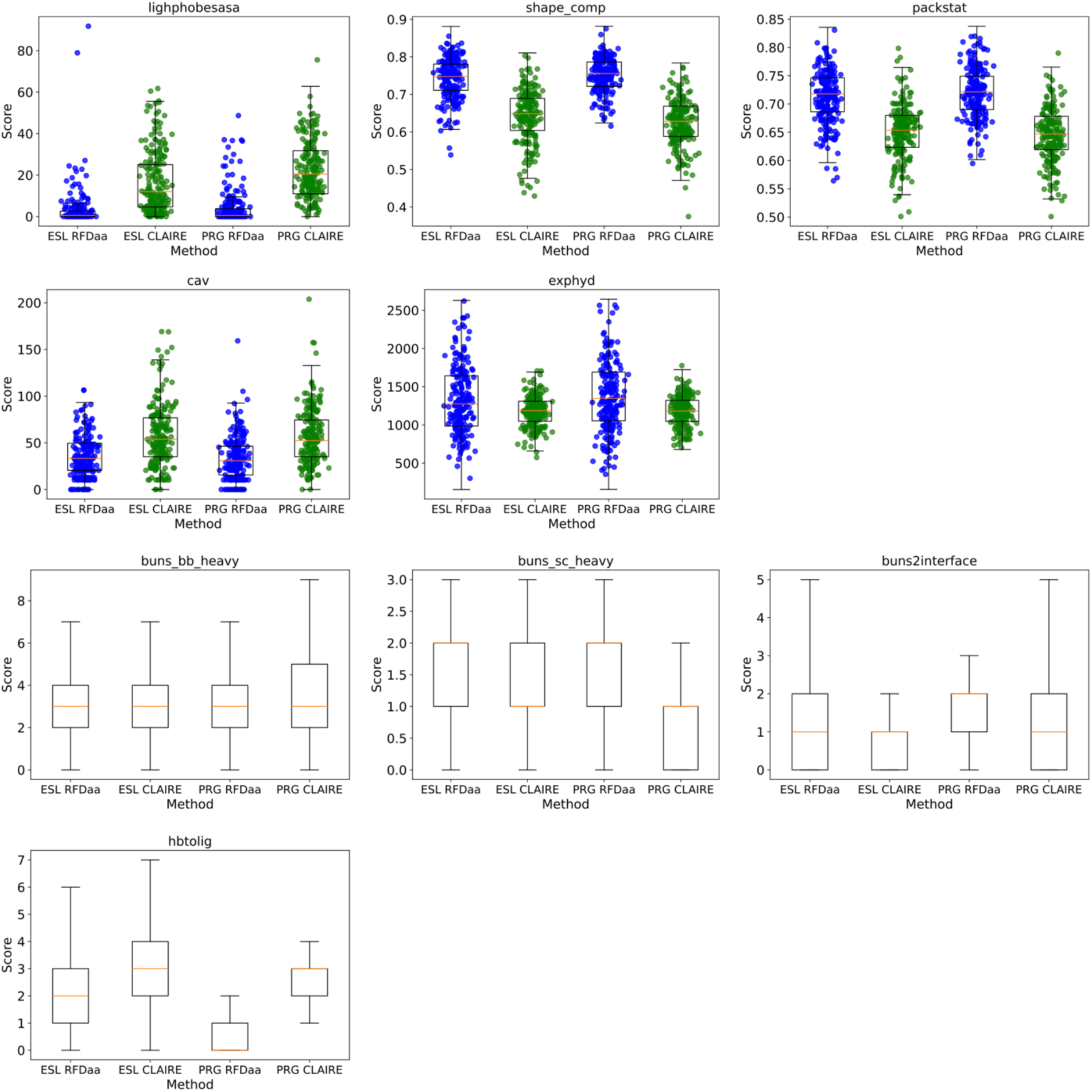
Boxplots comparing score distributions for CLAIRE designs (green) and RFDaa designs (blue) across relevant metrics. For metrics details see **Supplementary Table S1**. Boxes span the first to third quartile, with orange lines indicating the median value of the distribution. Whiskers extend to the furthest data point within 1.5 times beyond the interquartile range. For continuous distributions, a random sample of 100 values is overlayed across the box and whisker plot.

**Supplementary Figure S7:**
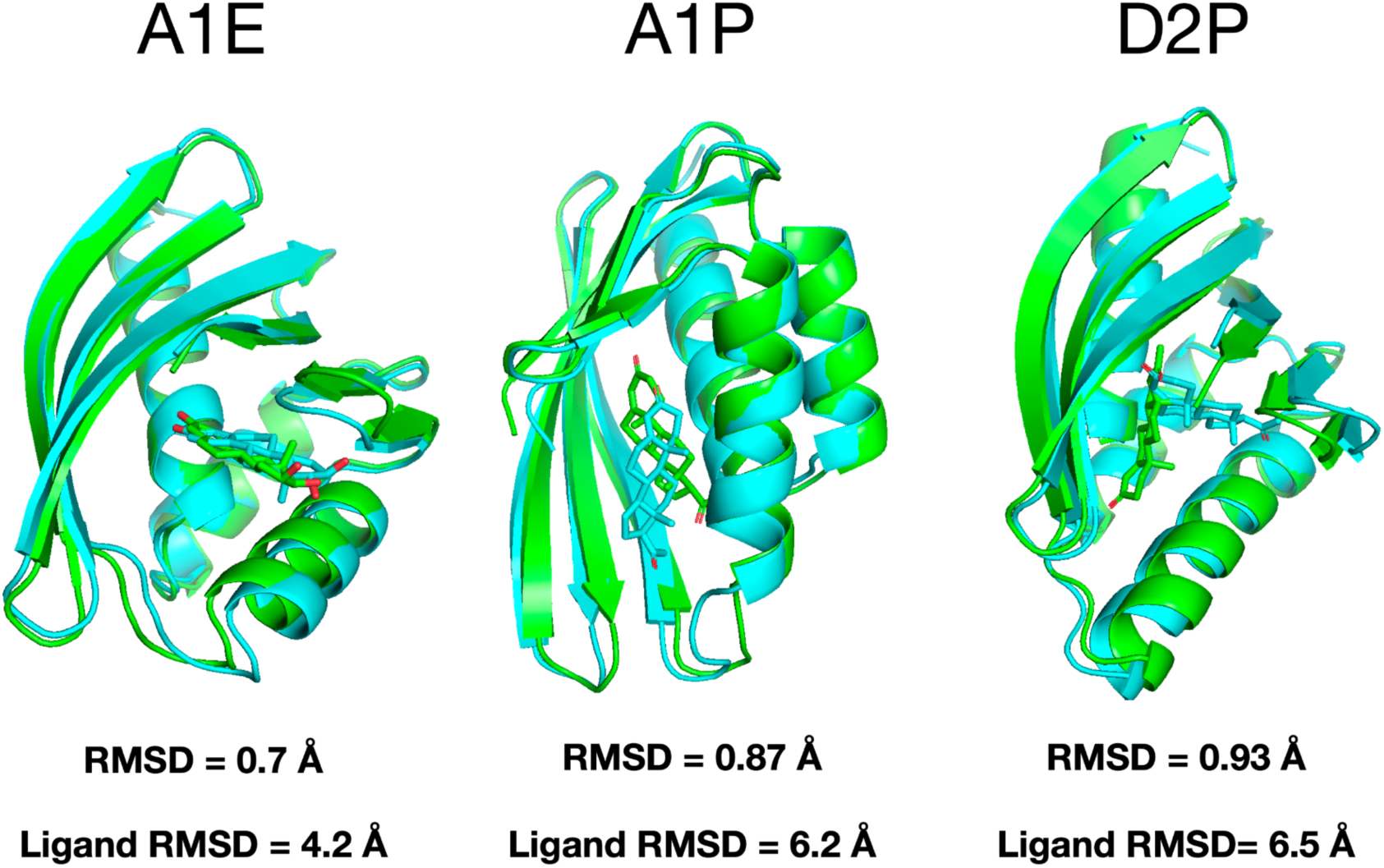
Alignment between design model and boltz-2 prediction for three experimentally characterized binders. Design models are shown in cyan and boltz-2 predictions are shown in green. The alpha carbon RMSD across all residues is given, along with the RMSD between the ligands in the alignment shown. While the protein structures are in good agreement, the ligands are not, being twisted and displaced in the predicted complex relative to the design models.

**Supplementary Figure S8:**
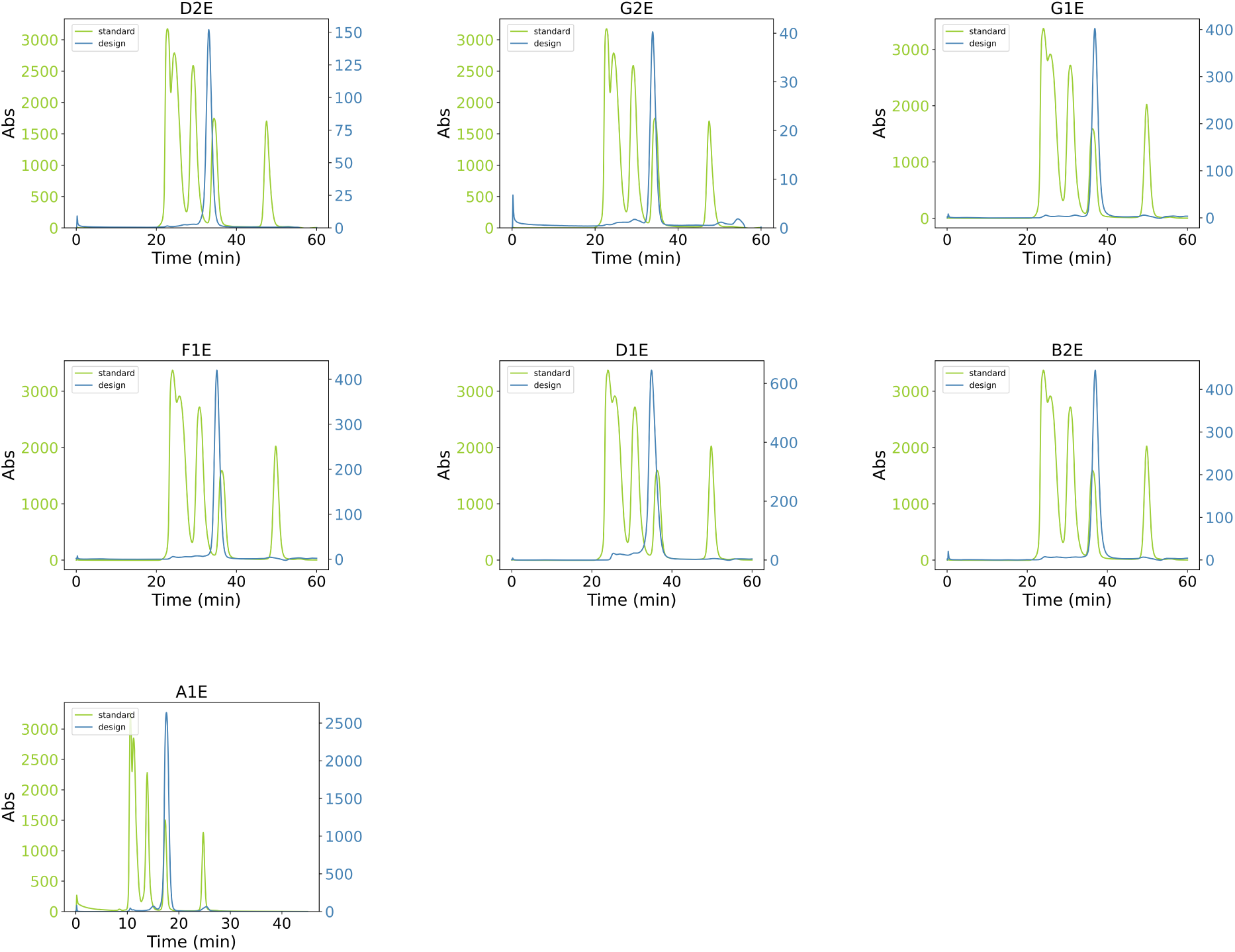
Size exclusion chromatography results for estriol binder designs. The green line corresponds to the standard, with the weight of the proteins for each peak being, from left to right, 670 kDa, 158 kDa, 44 kDa, 17 kDa, and 1.35 kDa. The blue lines correspond to our designed proteins, for which the molecular weights vary between 13-15 kDa.

**Supplementary Figure S9:**
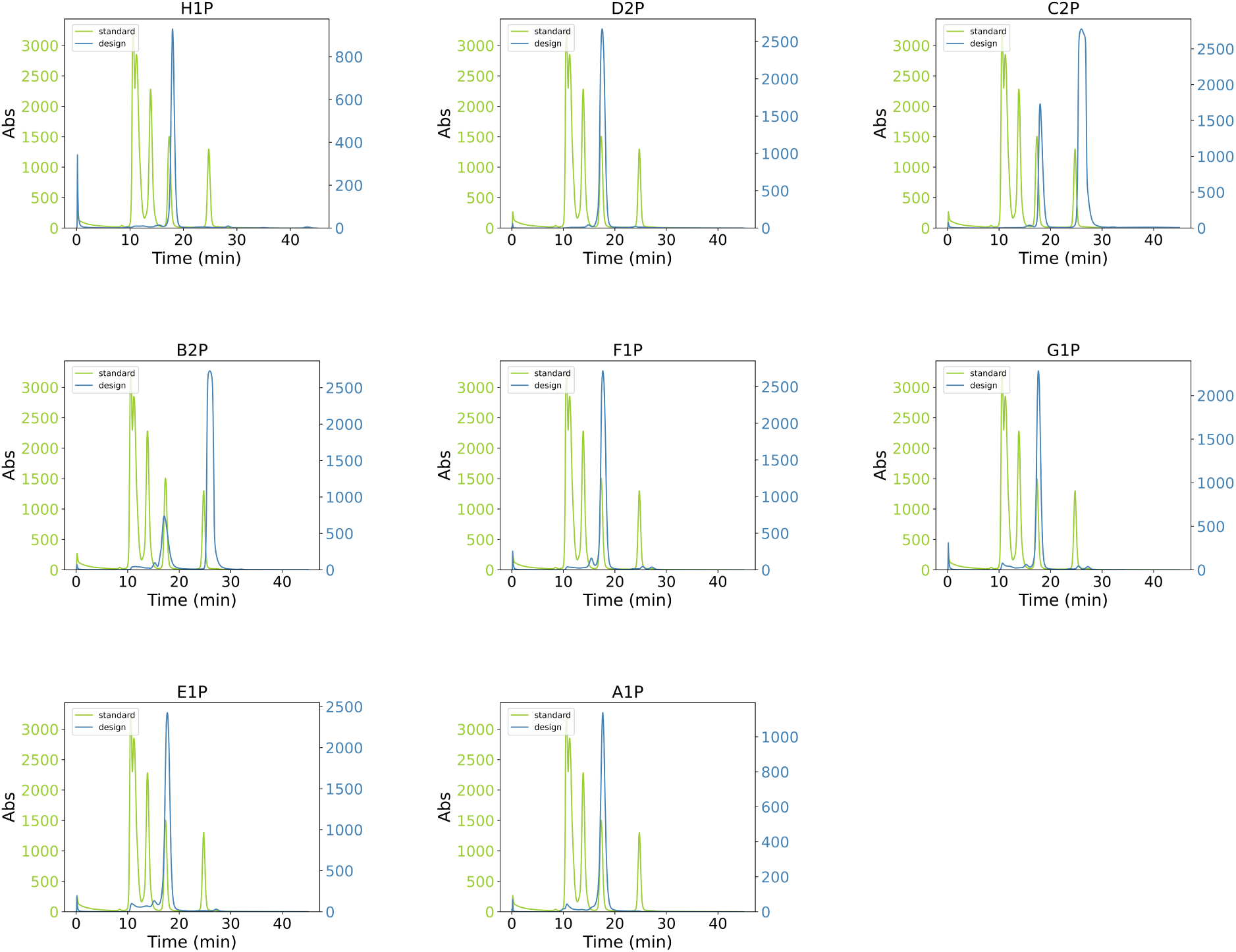
Size exclusion chromatography results for progesterone binder designs. The green line corresponds to the standard, with the weight of the proteins for each peak being, from left to right, 670 kDa, 158 kDa, 44 kDa, 17 kDa, and 1.35 kDa. The blue lines correspond to our designed proteins, for which the molecular weights vary between 13-15 kDa. Chromatograms for designs B2P and C2P were taken after titration with progesterone, which is responsible for the second peak at ∼28 minutes elution time.

**Supplementary Figure S10:**
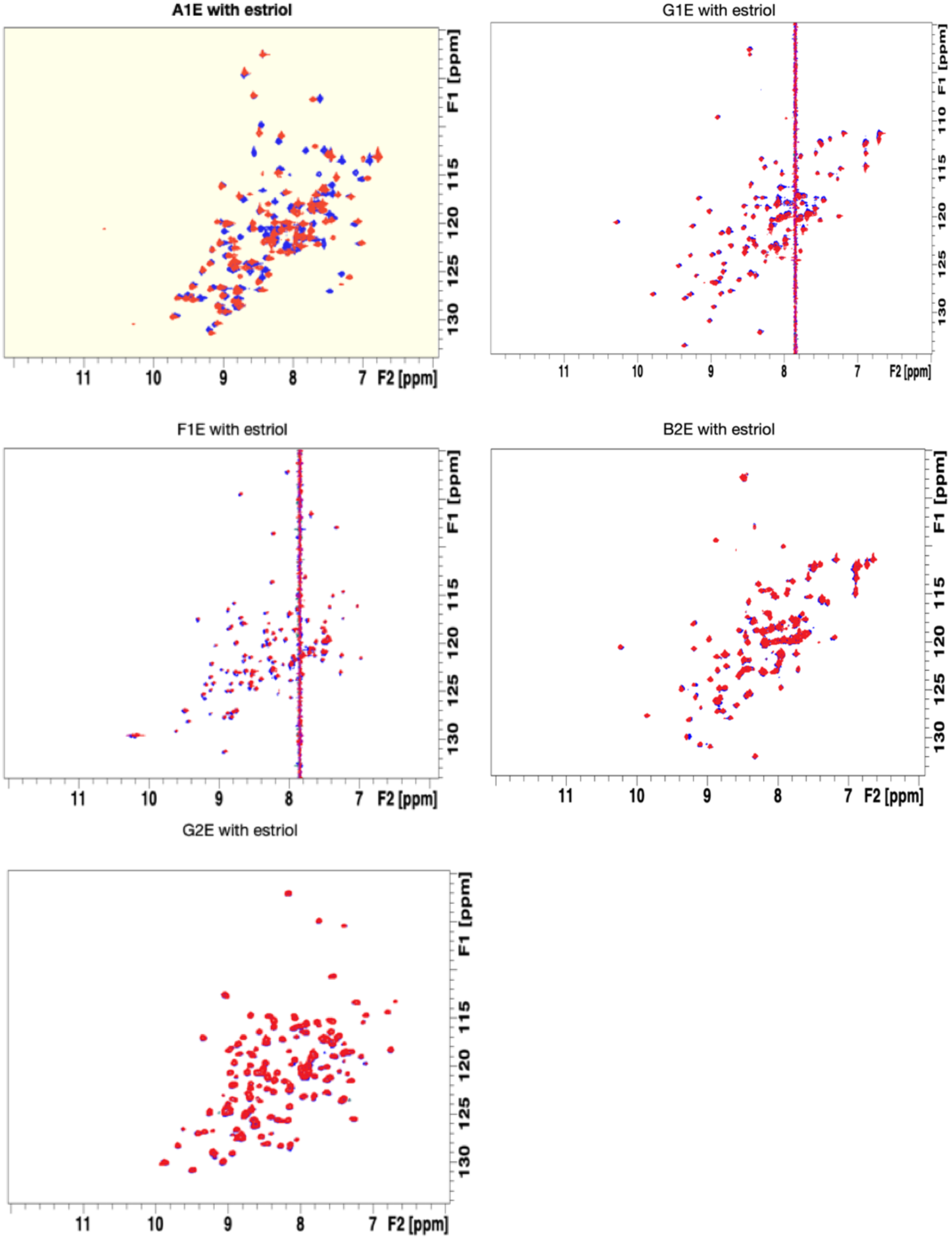
^15^N HSQC NMR chemical shift perturbations for designed estriol binders (only those folded by NMR are shown) with target ligand estriol. Apo spectra are shown in blue and red spectra are taken in the presence of estriol. Bold titles and yellow shading indicate significant ligand induced chemical shifts. Vertical bands are the result of using non-deuterated organic co-solvents. Design A1E shows binding. Samples for A1E, F1E, G1E, and B2E contained 653 μM estriol in 5% dimethylformamide in PBS at pH 7.4, with protein concentrations of 376, 88, 102, and 376 μM, respectively. The G2E sample contained 34 μM estriol in 2% DMSO in PBS at pH 7.4 with a protein concentration of 96 μM.

**Supplementary Figure S11:**
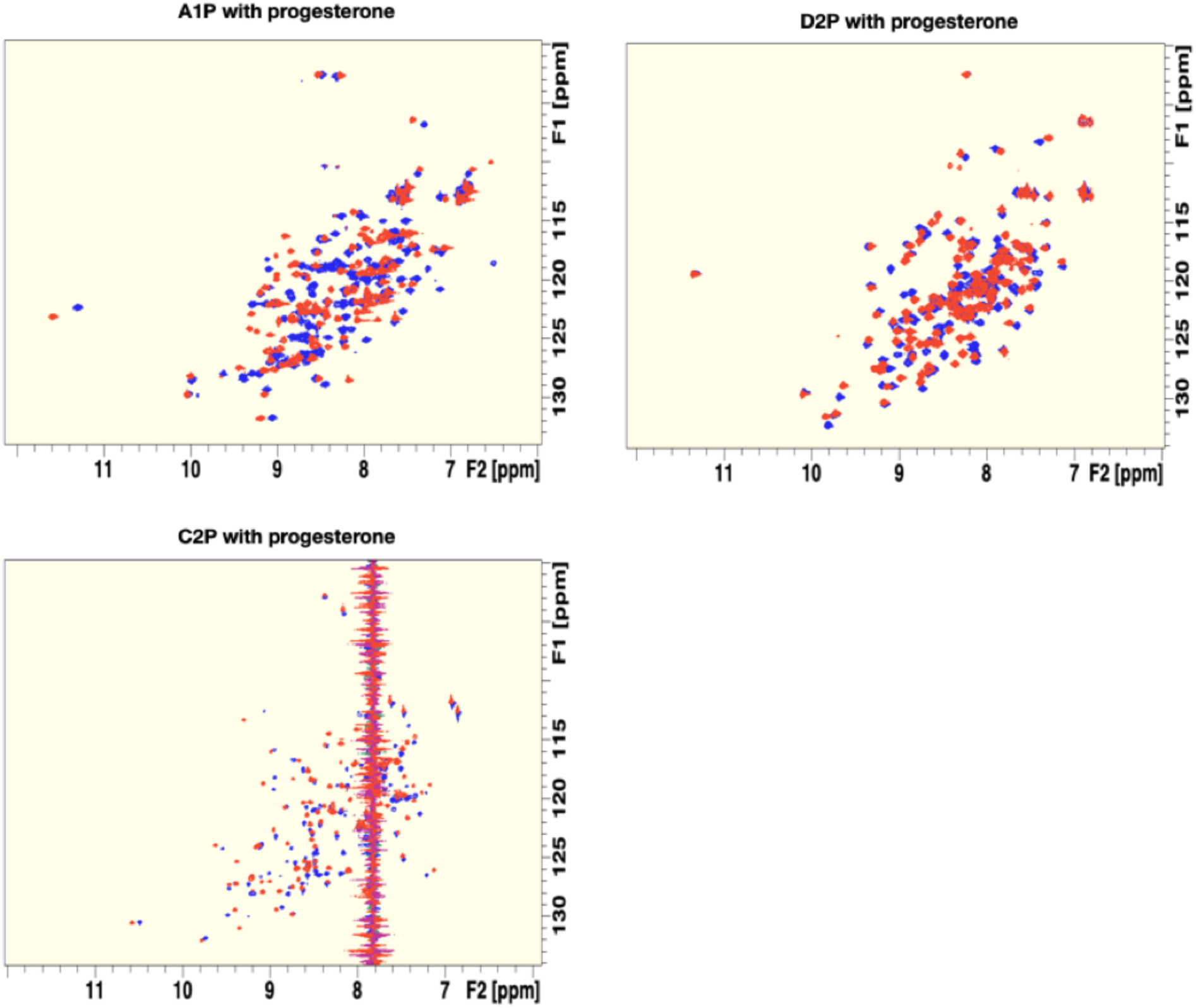
^15^N HSQC NMR chemical shift perturbations for designed progesterone binders (only those folded by NMR are shown) with target ligand progesterone. Apo spectra are shown in blue and red spectra are taken in the presence of progesterone. Bold titles and yellow shading indicate significant ligand induced chemical shifts. Vertical bands are the result of using non-deuterated organic co-solvents. All 3 of the designed progesterone binders show binding. A1P, C2P, and D2P samples contained 1.06 mM progesterone in 5% dimethylformamide in PBS at pH 7.4, with protein concentrations of 384, 74, and 466 μM, respectively.

**Supplementary Figure S12:**
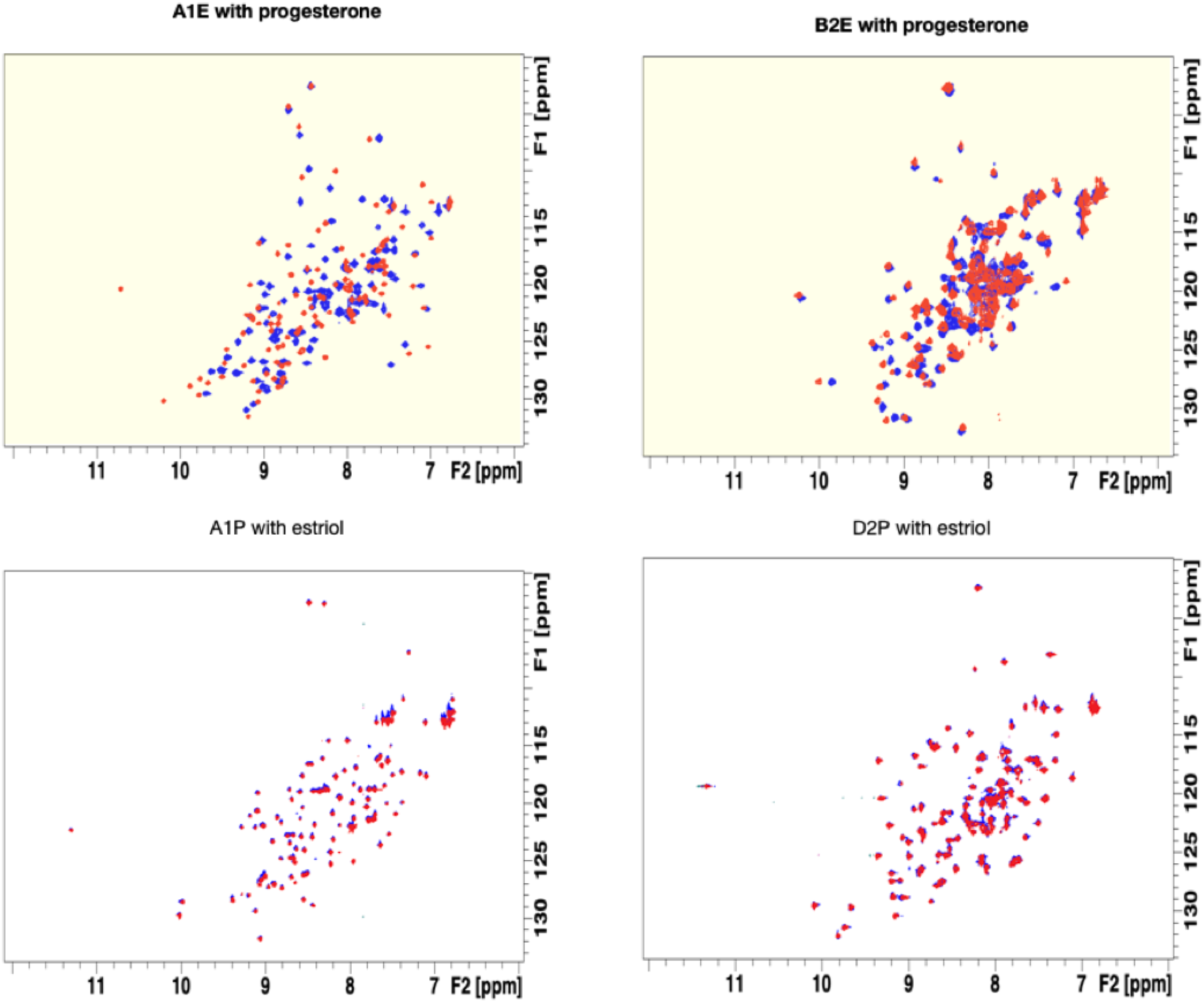
**Promiscuity of identified binders for off-target steroid**. Apo ^15^N HSQC NMR spectra are shown in blue and red. ^5^N HSQC NMR spectra are taken in the presence of ligand. Bold titles indicate significant ligand induced chemical shifts. The estriol binders A1E and B2E show strong chemical shift changes with progesterone, while the progesterone binders A1P and D2P do not show chemical shift changes with estriol. This behavior can be rationalized by the additional hydroxyl group in estriol relative to progesterone, which the progesterone binders are not designed to hydrogen bond with (see **Supplementary Figure S13**). Samples for A1E and B2E contained 363 μM progesterone in 5% dimethylformamide in PBS at pH 7.4, with protein concentrations at 376 μM. For A1P and D2P, samples contained 414 μM estriol in sodium phosphate buffer at pH 7.4, with protein concentrations of 93 μM and 120 μM, respectively.

**Supplementary Figure S13:**
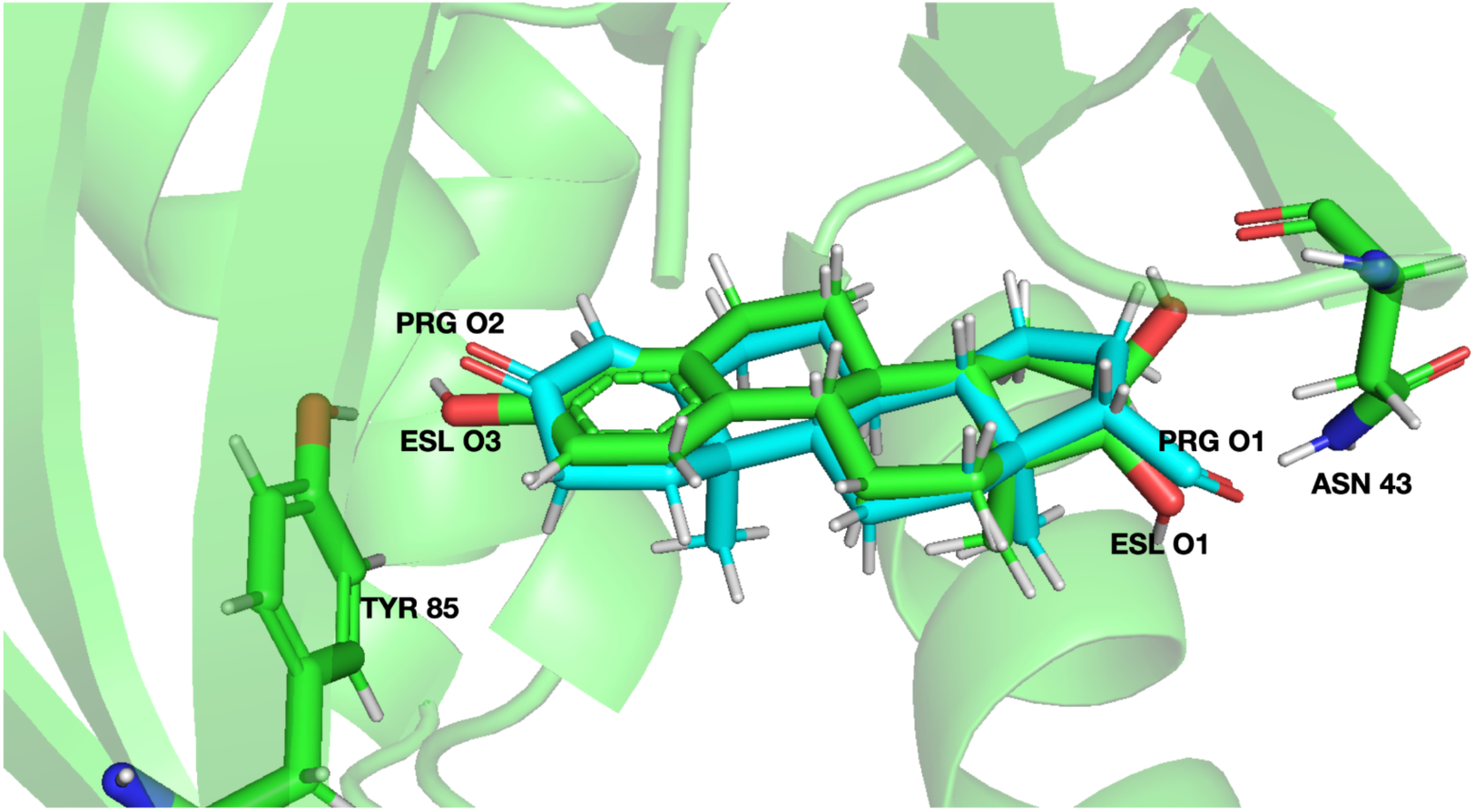
Overlay of estriol and progesterone modeled within the binding site of designed estriol binder A1E. The estriol is shown in green, while the progesterone is cyan. The proximity of the carbonyl oxygens in progesterone (prg-O2, prg-O1) to the equivalent hydroxyl oxygens (esl-O3, esl-O1) in estriol is apparent. Two residues, TYR-85 and ASN-43, which donate hydrogen bonds to these estriol oxygens in the A1E design model, appear capable of donating hydrogen bonds to the progesterone oxygens with minimal conformational adjustment. Progesterone was modeled in the binding site by manually mapping its carbons to the equivalent positions in the estriol steroid backbone using the Pymol pair_fit command.

**Supplementary Figure S14:**
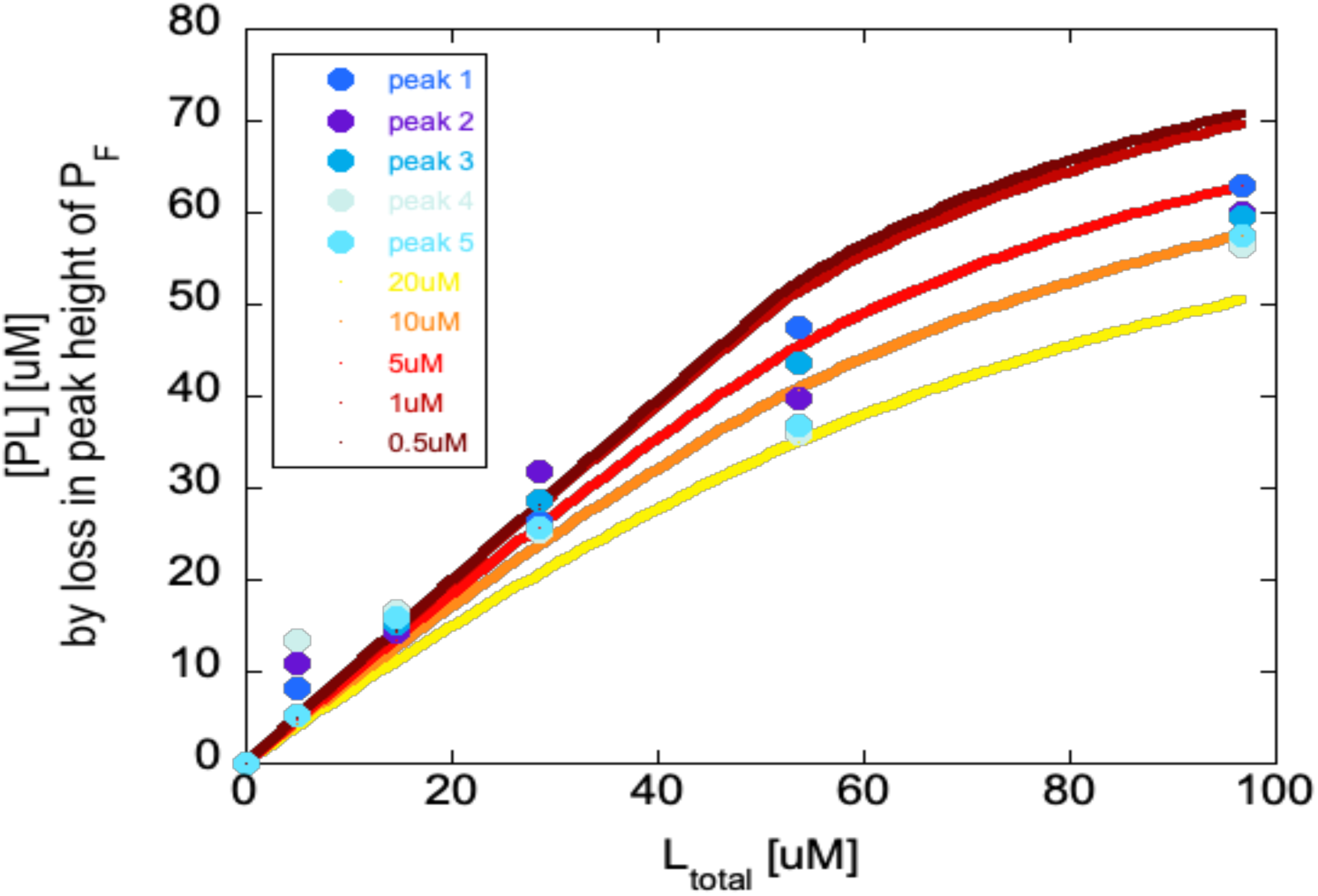
NMR titrations for D2P with progesterone. Addition of low micromolar concentrations of progesterone leads to successive disappearance of protein peaks in the ^15^N HSQC NMR spectra, consistent with progesterone binding in the intermediate exchange regime. Under the assumption that loss in peak intensity can be used to estimate the fraction of progesterone bound protein, we obtained the titration curves shown in the figure for several well-separated peaks (filled circles in shades of blue). As the protein concentration in this experiment is high (around 70 μM) compared to the dissociation constant *K_d_*, the curves cannot be fit reliably. However, overlay of simulated curves using the quadratic binding equation (AB- complex between ligand A and protein B) 

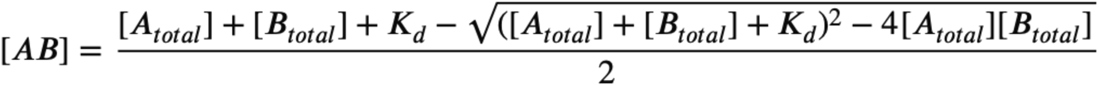

and assuming Kd values between 0.5 and 20 μM estimates a *K_d_* between 5 μM (bright red curve) and 10 μM (orange curve).

**Supplementary Figure S15:**
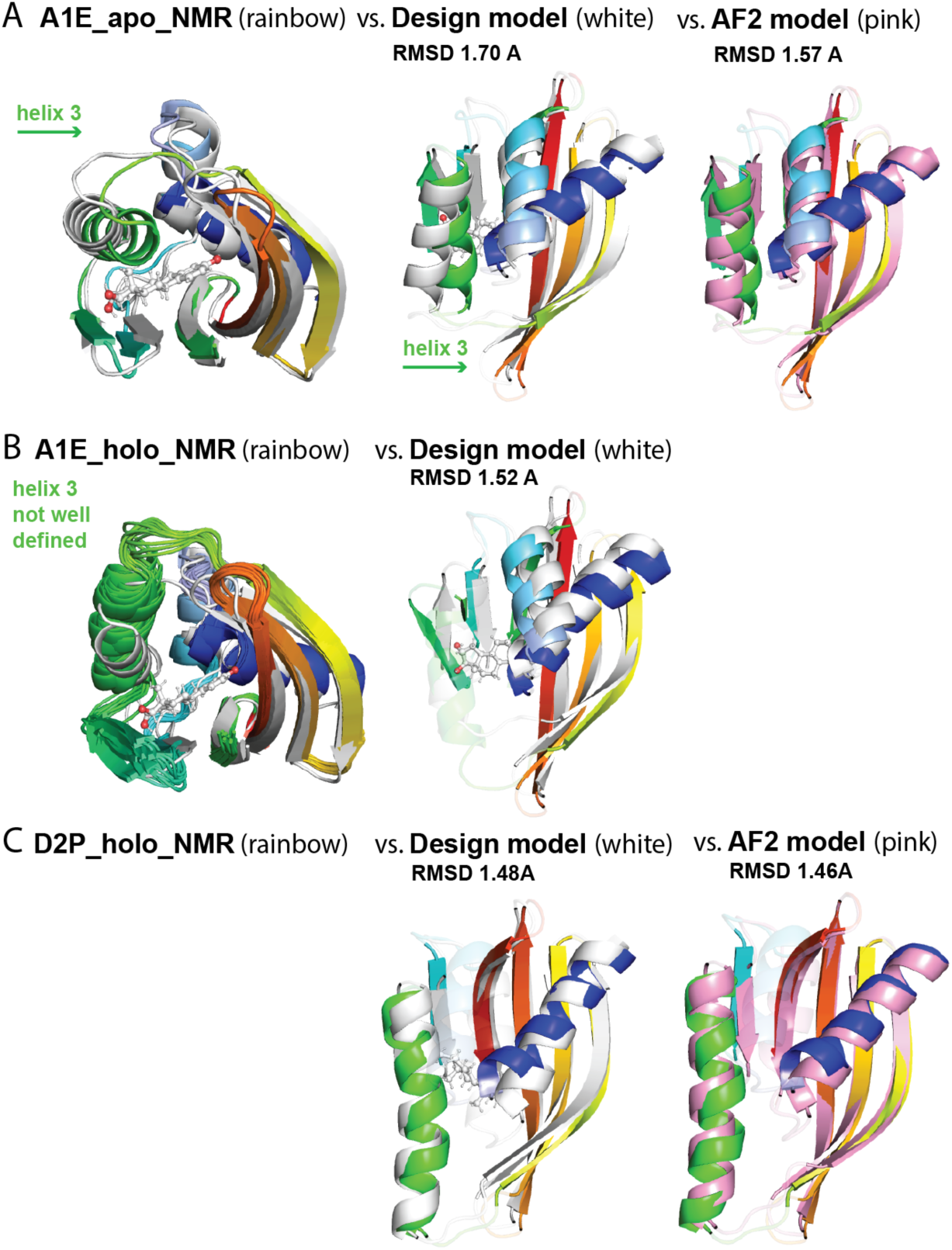
Structural details comparing NMR structures, AF2 models, and design models. **A**) NMR structure (rainbow) for design A1E in the apo state overlaid with the design model (white) shows a change in helix 3 that moves into the (empty) binding pocket in the NMR structure. This helix shift is better predicted in the AF2 model (pink, right plot, also shown in main Fig. 4) compared to the design model shown in the same orientation (middle plot, white). **B**) NMR model of design A1E (rainbow) with progesterone added overlayed with the design model (white) shows that helix 3 (green) is not well defined. The right plot shows the comparison between the NMR structure and the design model omitting helix 3 and loop regions. **C**) The NMR model for design D2P in the holo state is predicted well by both the design model (white, left) and the AF2 model (pink, right, also shown in main Figure 4). Regions used in the RMSD calculation are colored in all panels.

**Supplementary Figure S16:**
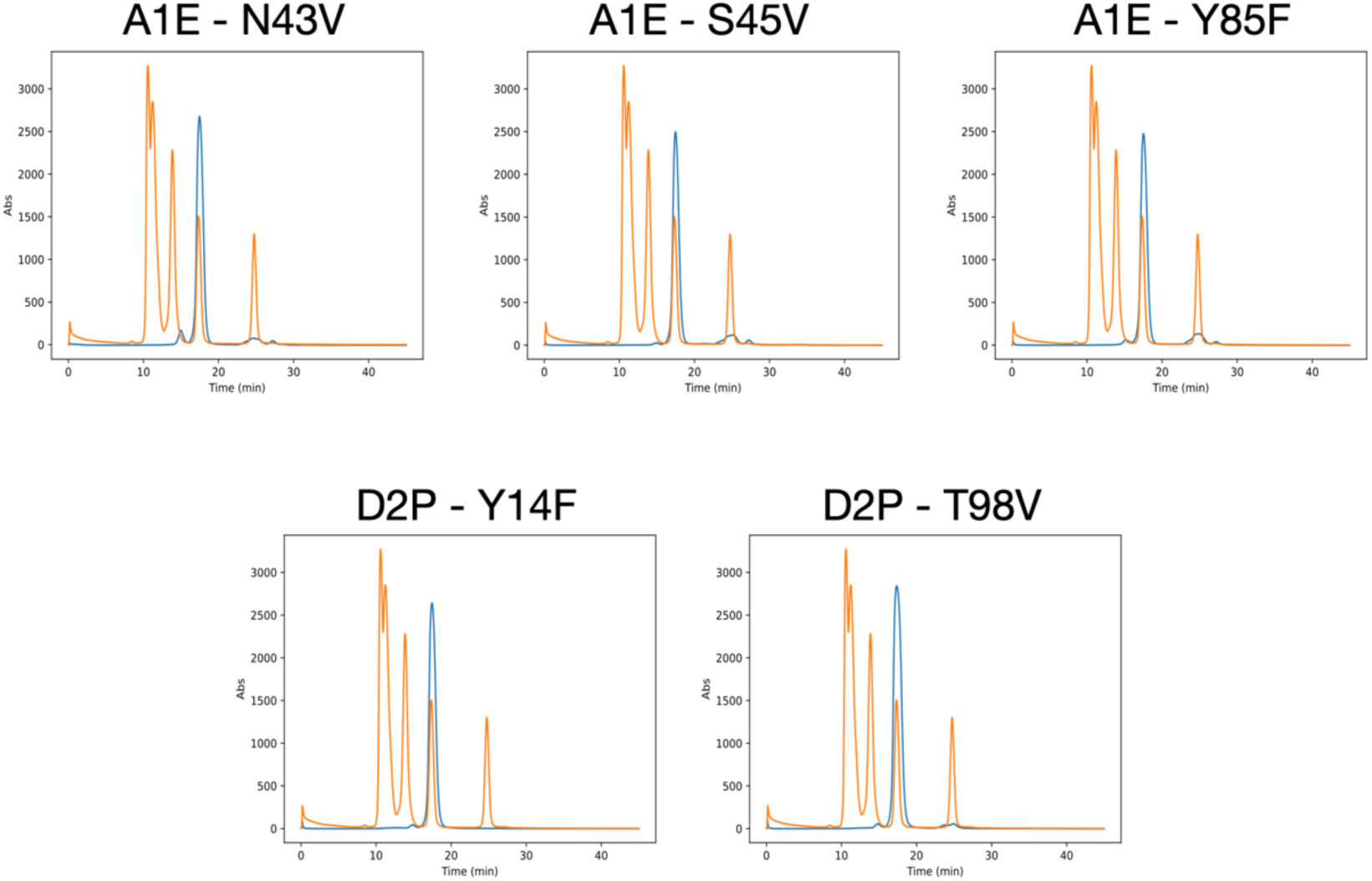
Size exclusion chromatography traces for point mutants. The orange line corresponds to the standard, with the weight of the proteins for each peak being, from left to right, 670 kDa, 158 kDa, 44 kDa 17 kDa, and 1.35 kDa. The blue lines correspond to the point mutants, for which the molecular weights vary between 13-15 kDa. The source design along with the mutation made are indicated above each chromatogram.

**Supplementary Figure S17:**
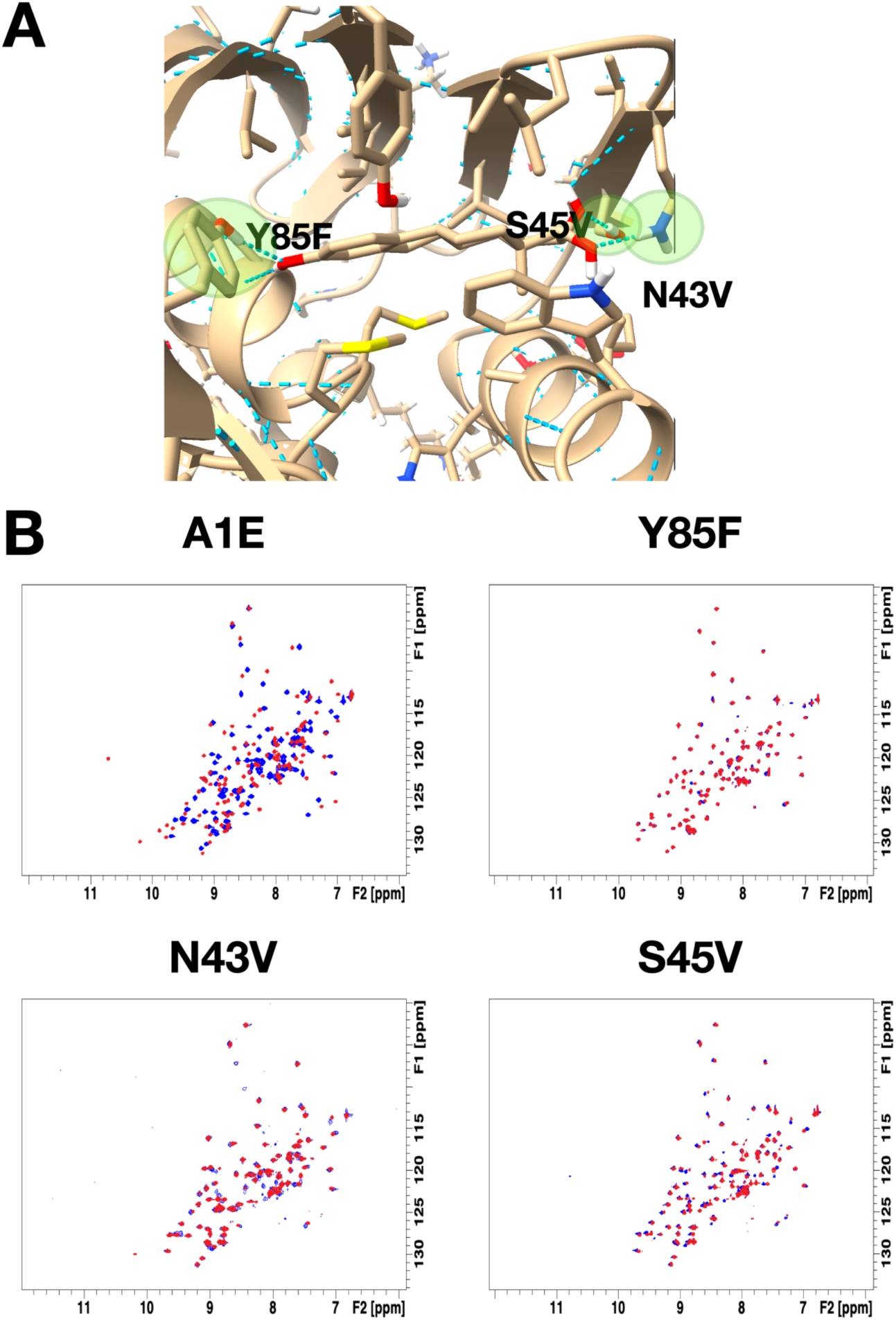
^15^N HSQC NMR chemical shift perturbation experiments on A1E point mutants targeting key protein-ligand hydrogen bonds. **A)** Close up of A1E binding site, highlighting residues targeted for mutation and indicating the substitution made. **B)** ^15^N HSQC NMR chemical shift perturbations for original design A1E (top left, as in Figure 3D) and 3 point mutants. In all cases, the apo spectra are shown in blue, and the red spectra are taken with progesterone in solution. Samples for the Y85F, N43V, and S45V mutants contained 1.5 mM progesterone in 5% dimethylformamide in PBS at pH 7.4, with protein concentrations of 274 μM, 313 μM, and 260 μM, respectively.

**Supplementary Figure S18:**
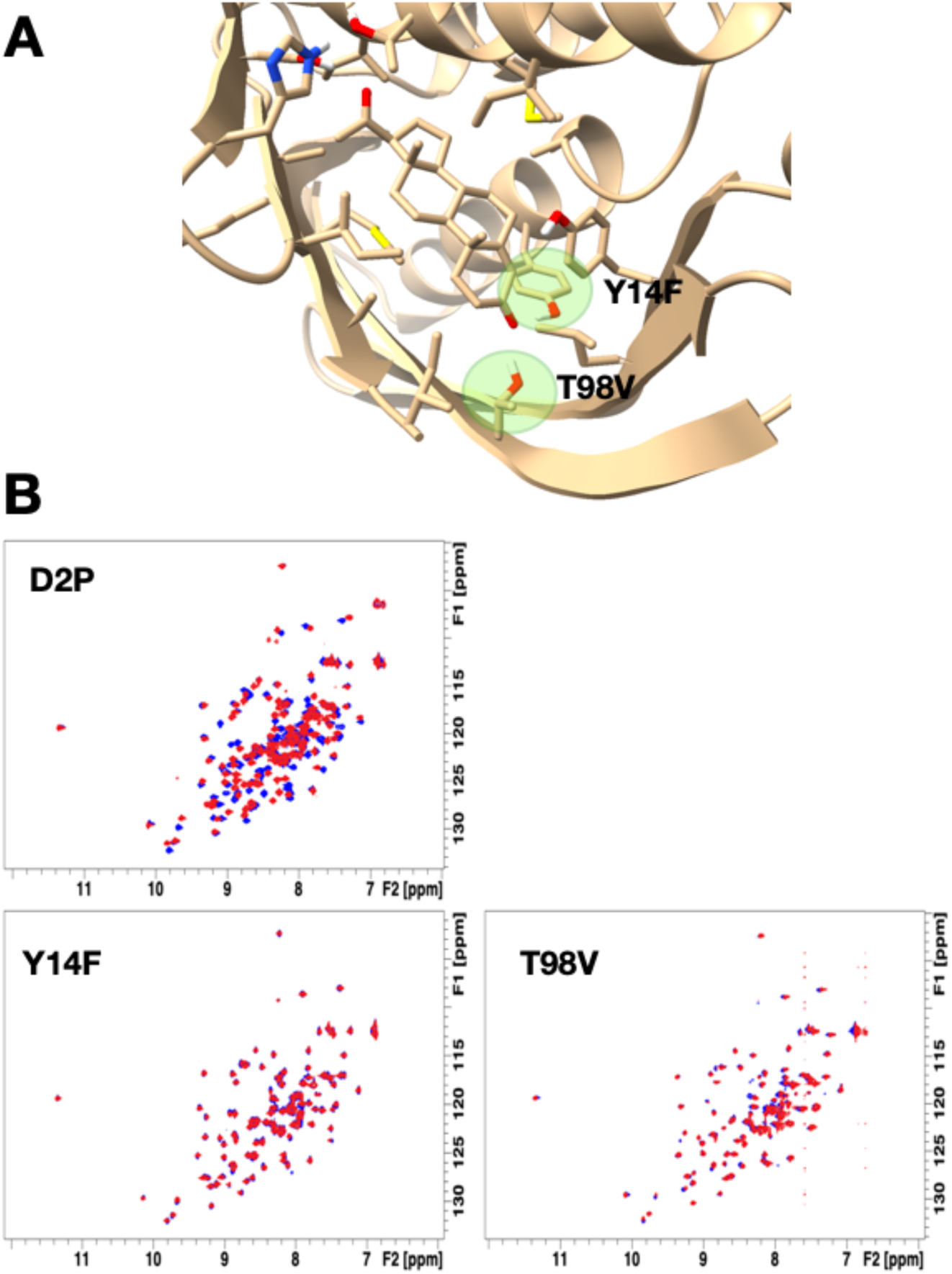
^15^N HSQC NMR chemical shift perturbation experiments on D2P point mutants targeting key protein-ligand hydrogen bonds. **A)** Close up of D2P binding site, highlighting residues targeted for mutation and indicating the substitution made. **B) ^1^**^5^N HSQC NMR chemical shift perturbations for original design D2P (top left, as in Figure 3D) and 2 point mutants. In all cases, the apo spectra are shown in blue, and the red spectra are taken with progesterone in solution. Samples for the Y14F and T98V mutants contained 1.5 mM progesterone in 5% dimethylformamide in PBS at pH 7.4, with protein concentrations of 346 μM and 317 μM, respectively.

**Supplementary Figure S19:**
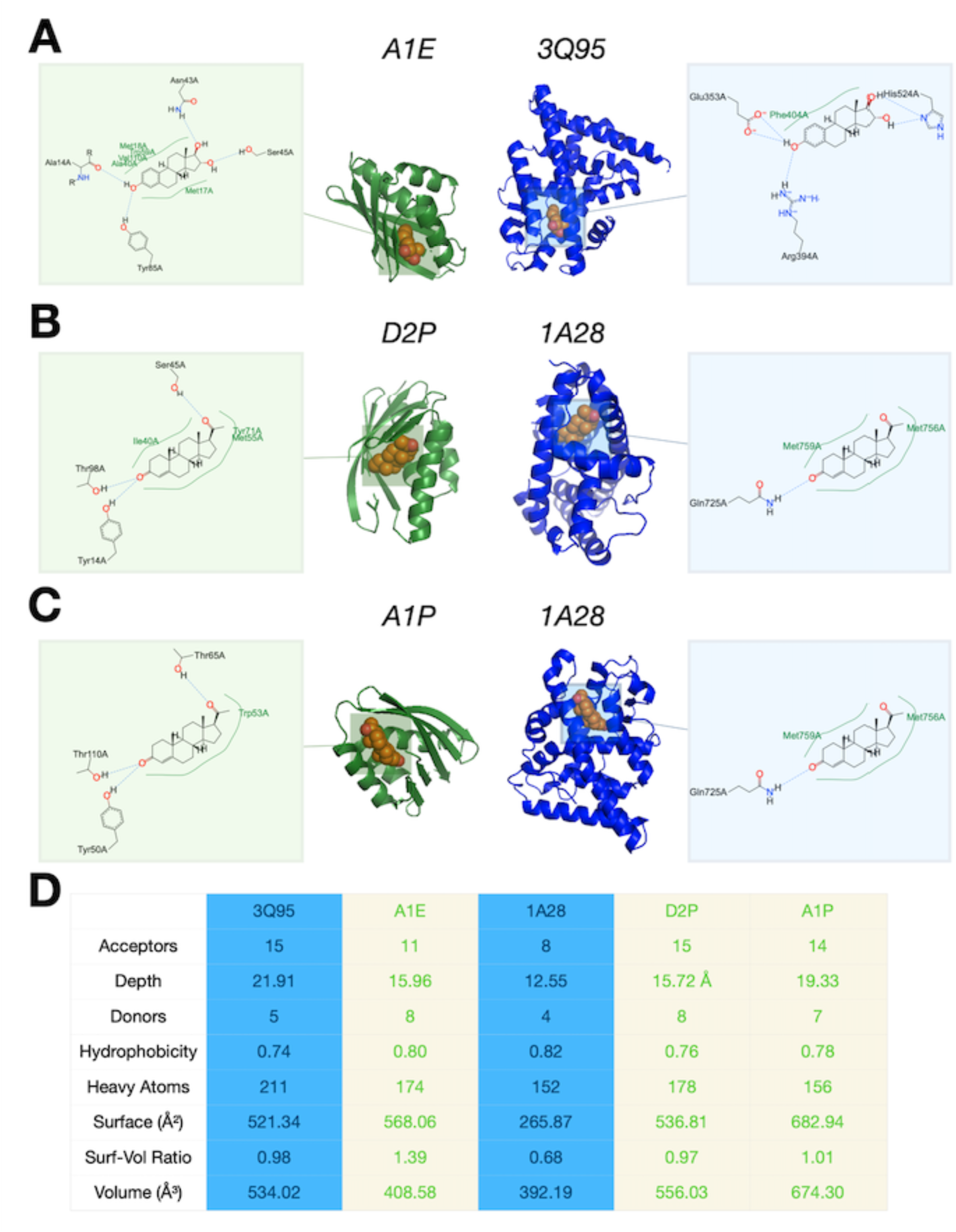
Comparison of binding site interactions in designed estriol and progesterone binders with binding site interactions in human estrogen receptor. **A)** Comparison between designed estriol binder A1E (green) and human estrogen receptor with estriol bound (blue, PDB 3Q95). Schematics show binding site interactions, with protein-ligand hydrogen bonds indicated by dashed lines and protein residues that pack with ligand indicated by green text. **B)** Comparison between designed progesterone binder D2P (green) and human estrogen receptor with progesterone bound (blue, PDB 1A28). **C)** Comparison between designed progesterone binder A1P (green) and human estrogen receptor with progesterone bound (blue, PDB 1A28). **D)** Table showing binding pocket properties for designed binders (green) and human estrogen receptor (blue). Pockets are defined using a grid-based approach (*53*) wherein grid points with no protein atoms within a distance equal to their Van der Waals radii are clustered and merged until they can no longer be expanded in any direction without including points occupied by protein atoms. The acceptor, donor, and heavy atom counts refer to the number of each respective atom that is considered part of the pocket (defined by the distance between the atom and any pocket grid point being less than the sum of the atoms Van der Waals radius, the grid spacing factor, and the cubic grid diagonal). Depth (Å) and volume (Å^3^) are calculated by summing values for individual grid points within the pocket, while surface (Å^2^) is calculated by summing the SASA values of all pocket atoms. Hydrophobicity is the fraction of pocket atoms that are hydrophobic.

**Supplementary Figure S20:**
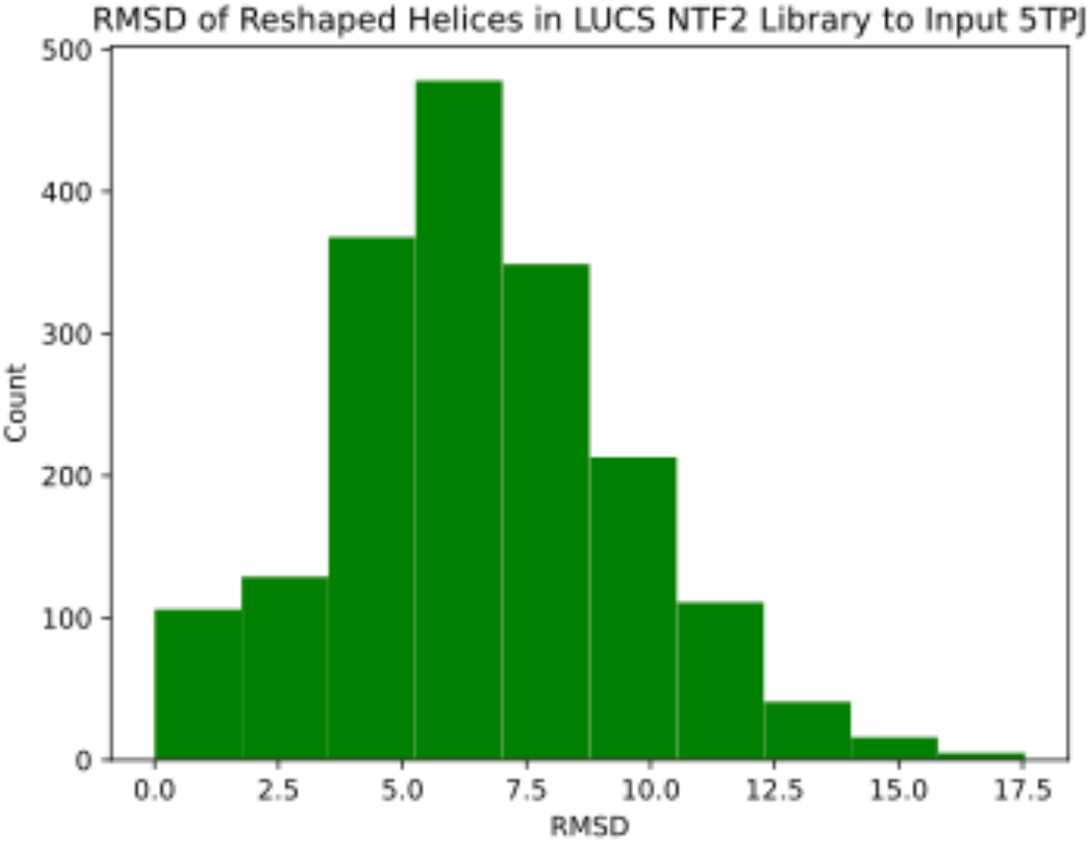
Distribution of RMSD values between the reshaped helices in the LUCS NTF2 scaffold library and those of the original input, PDB 5TPJ.

**Supplementary Table S1:** Glossary of filtering criteria employed throughout design workflow. Metric names are given as they appear in plots for score distributions, along with the values used for filtering thresholds and a brief description of what each term measures.

| Metric | Filter Threshold | Significance |
| --- | --- | --- |
| boltz | $\leq -0.3$ | Average Boltzmann probability of binding site residue rotamers |
| bsE_per_res (REU) | $\leq -2.0$ | Rosetta energy per binding site residue |
| buns_bb | $\leq 5$ | Buried unsatisfied backbone polar atoms |
| buns_sc | $\leq 1$ | Buried unsatisfied side chain polar atoms |
| buns2interface | 0 | Number of buried unsatisfied polar atoms at interface |
| Ca RMSD (Å) | $\leq 1.5$ | Similarity design model:ColabFold prediction |
| cav (Å<sup>3</sup>) | $\leq 120$ | Cavity volume (with ligand bound) |
| contact_molsurf (Å<sup>2</sup>) | $\geq 151$ | Contact molecular surface |
| ddg | $\leq -23$ | Difference in Rosetta energy between complex and components |
| dsasa | $\geq 0.7$ | Fraction ligand SASA buried upon binding |
| exphyd | $\leq 1200$ | Exposed hydrophobic surface area of protein |
| Hb2lig | $\geq 3$ | Number of protein-ligand hydrogen bonds |
| lig_hphobe_sasa (Å<sup>2</sup>) | $\leq 40.0$ | Exposed ligand hydrophobic SASA |
| lig_oversat | 0 | Oversaturated polar atoms on ligand |
| oversat | 0 | Oversaturated polar atoms across protein |
| packstat | $\geq 0.65$ | Rosetta packing statistic, measure of packing efficiency |
| pLDDT | $\geq 88.0$ | ColabFold confidence metric |
| shape_comp | $\geq 0.6$ | Protein:ligand shape complementarity |

**Supplementary Table S2:**
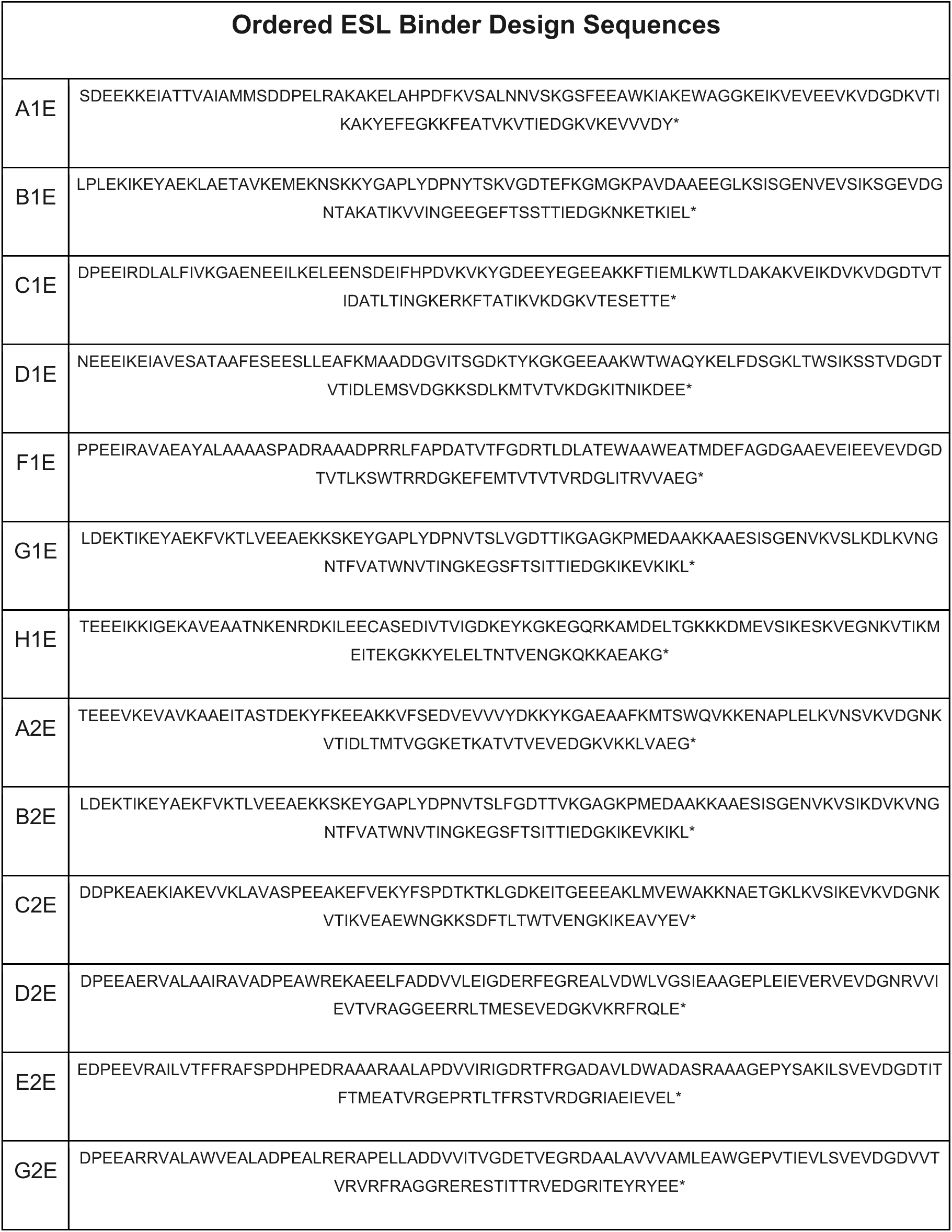
Amino acid sequences for experimentally characterized estriol binder designs.

**Supplementary Table S3:**
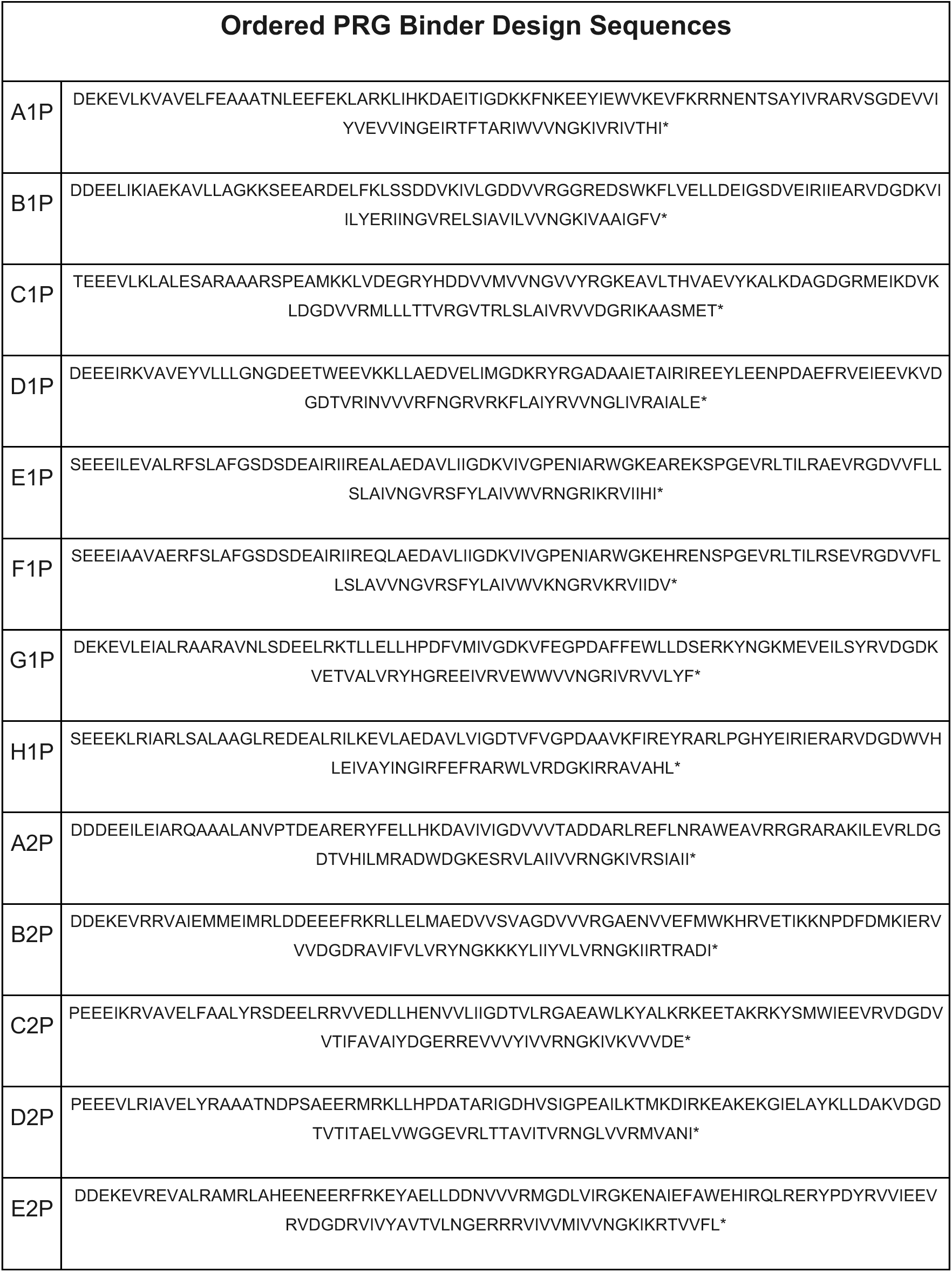
Amino acid sequences for experimentally characterized progesterone binder designs.

**Supplementary Table S4:**
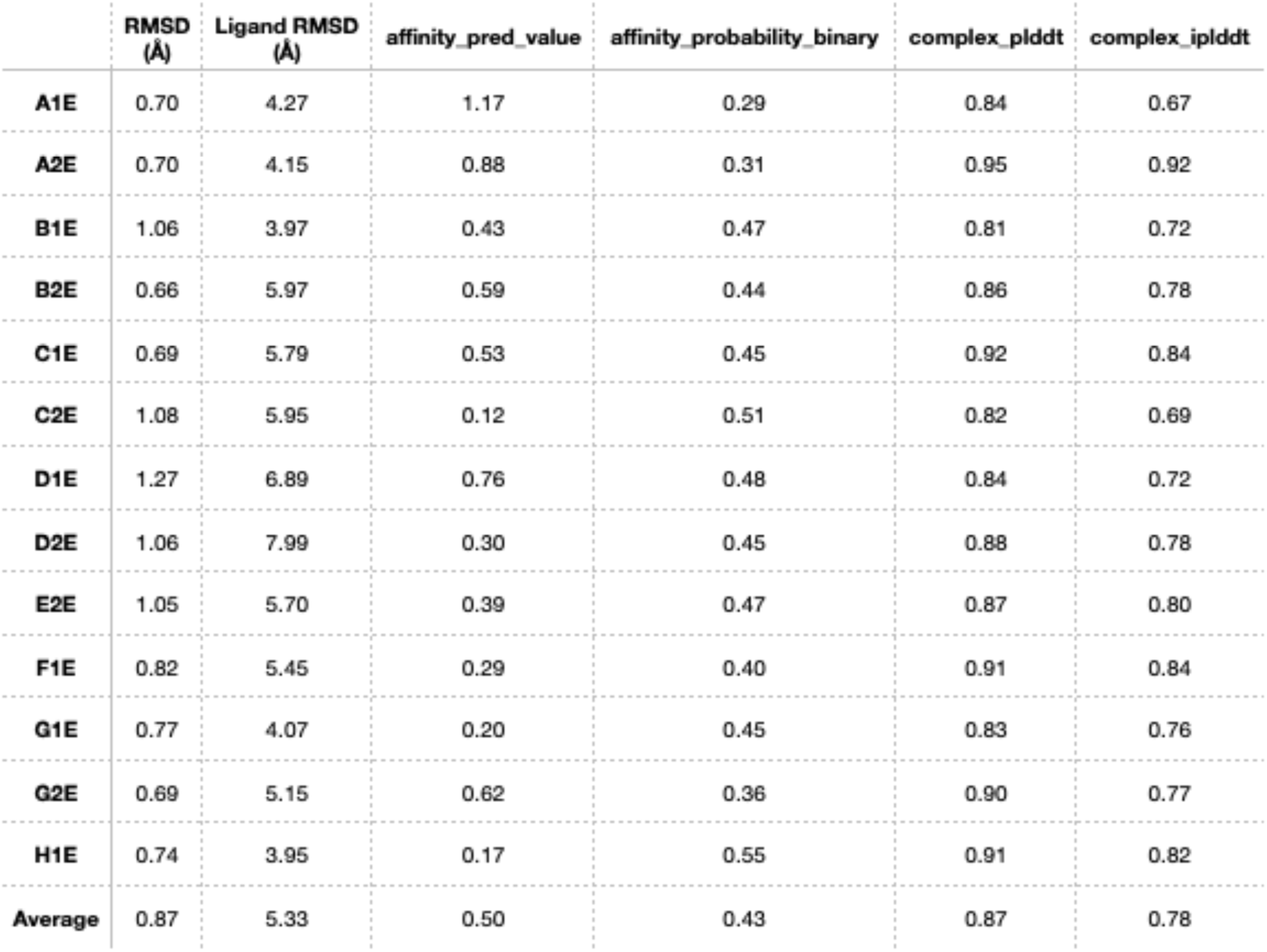
Results of Boltz-2 predictions for experimentally characterized estriol binder designs. RMSD values are for alpha carbons across the full proteins, while ligand RMSD is across all ligand atoms after aligning the proteins so as to minimize their global Ca RMSD. The affinity_pred_value is the predicted - log(IC50) for the complex, with higher values indicating higher predicted affinity. The affinity_probability_binary is the predicted probability that the target molecule binds the protein. The complex_plddt is the model’s confidence for the predicted structure, while the iplddt is the confidence for the predicted interface.

**Supplementary Table S5:**
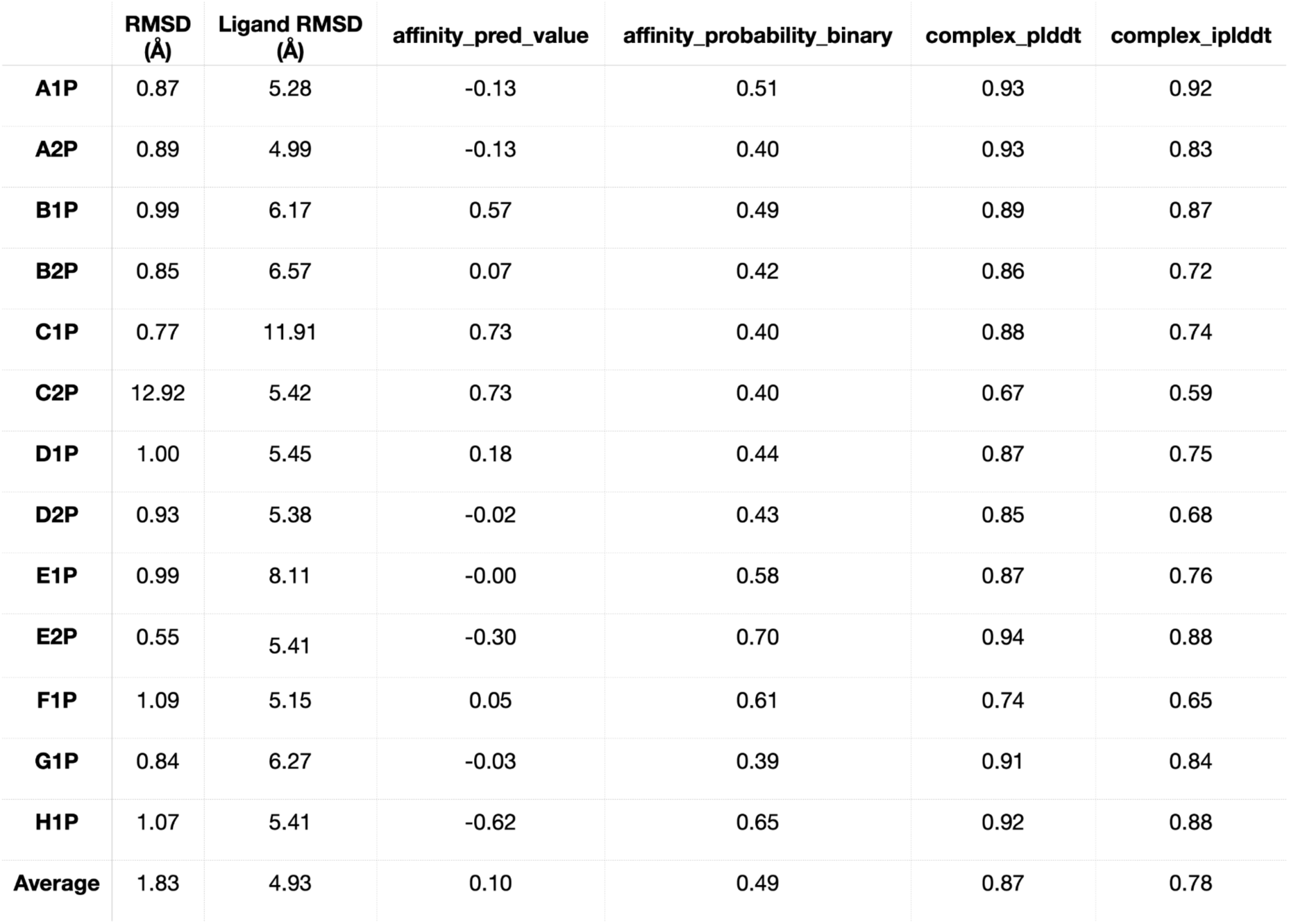
Results of Boltz-2 predictions for experimentally characterized progesterone binder designs. RMSD values are for alpha carbons across the full proteins, while ligand RMSD is across all ligand atoms after aligning the proteins so as to minimize their global Ca RMSD. The affinity_pred_value is the predicted -log(IC50) for the complex, with higher values indicating higher predicted affinity. The affinity_probability_binary is the predicted probability that the target molecule binds the protein. The complex_plddt is the model’s confidence for the predicted structure, while the iplddt is the confidence for the predicted interface.

**Supplementary Table S6.** Experimental success rates. Y: yes; N: no; ND: not determined.

| Design | monomeric | folded (NMR) | binds ESL (CSP) | binds PRG (CSP) |
| --- | --- | --- | --- | --- |
| **Estriol binder designs** |  |  |  |  |
| A1E | Y | Y | Y | Y |
| B1E | N |  |  |  |
| C1E | N |  |  |  |
| D1E | Y | N |  |  |
| F1E | Y | Y | N | ND |
| G1E | Y | Y | N | ND |
| H1E | N |  |  |  |
| A2E | N |  |  |  |
| B2E | Y | Y | N | Y |
| C2E | N |  |  |  |
| D2E | Y | N |  |  |
| E2E | N |  |  |  |
| G2E | Y | Y | N | ND |
| NUMBER | 7/13 | 5/13 | 1/13 | 2/13 |
| **Progesterone binder designs** |  |  |  |  |
| A1P | Y | Y | N | Y |
| B1P |  |  |  |  |
| C1P |  |  |  |  |
| D1P |  |  |  |  |
| E1P | Y | N |  |  |
| F1P | Y | N |  |  |
| G1P | Y | N |  |  |
| H1P | Y | ND |  |  |
| A2P |  |  |  |  |
| B2P | Y | N |  |  |
| C2P | Y | Y | ND | Y |
| D2P | Y | Y | N | Y |
| E2P | ND | N |  |  |
| NUMBER | 8/13 | 3/13 | 0/13 | 3/13 |

**Supplementary Table S7:** Success rates from refinement steps in CLAIRE (shown in **Figure 1G**). 8,500 buried matches for each of estriol and progesterone were subject to binding site design using the Rosetta EnzymeDesign application followed by our HBRefine protocol (rows 3 and 4). For resultant designs passing all binding site metrics (260 and 578 for estriol and progesterone, respectively, row 2), ProteinMPNN was applied across non binding site residues, and the generated sequence profiles were fed to Rosetta FastDesign to produce ∼10 designs for each input structure. These design runs resulted in a total of 2,487 designed estriol binders and 5,111 designed progesterone binders, which were evaluated for both binding and stability metrics (row 5).

|  | PRG designs<br>passing filters | ESL designs<br>passing filters |
| --- | --- | --- |
| Rosetta Enzyme<br>Design; binding<br>metrics | 219/8,500 | 432/8,500 |
| HBRefined; binding<br>metrics | 260/8,500 | 578/8,500 |
| HBRefined; all<br>metrics | 80/8,500 | 82/8,500 |
| Stability refined; all<br>metrics | 130/2,487 | 346/5,111 |

**Supplementary Table S8:** Success rates from refinement steps in RFDiffusionAA (shown in **Figure 2A**). 6,000 backbones were generated around each of estriol and progesterone. For each backbone passing initial criteria (Methods), 3 rounds of design were performed using LigandMPNN followed by Rosetta FastRelax, producing 8 designs for each input, resulting in 31,653 designs for progesterone and 28,586 for estriol.

|  | PRG | ESL |
| --- | --- | --- |
| Backbone metrics | 3,968/6,000 | 3,585/6,000 |
| LMPNN; all metrics | 10/31,653 | 736/28,586 |
| HBRefined; all metrics | 30/31,653 | 933/28,586 |

**Supplementary Table S9:**
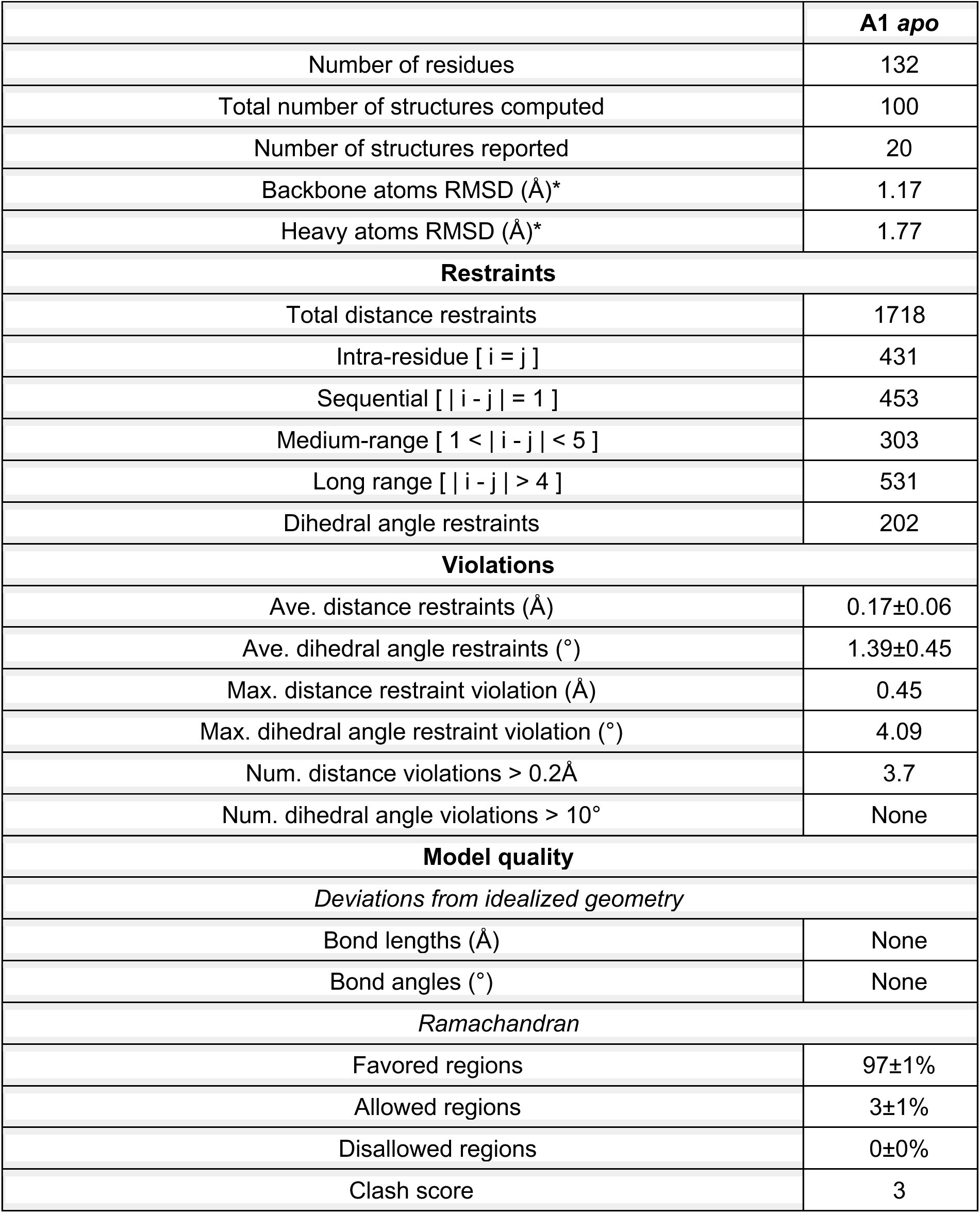
NMR structure statistics.

